# Emergent Speciation by Multiple Dobzhansky–Muller Incompatibilities

**DOI:** 10.1101/008268

**Authors:** Tiago Paixão, Kevin E. Bassler, Ricardo B. R. Azevedo

## Abstract

The Dobzhansky–Muller model posits that incompatibilities between alleles at different loci cause speciation. However, it is known that if the alleles involved in a Dobzhansky–Muller incompatibility (DMI) between two loci are neutral, the resulting reproductive isolation cannot be maintained in the presence of either mutation or gene flow. Here we show that speciation can emerge through the collective effects of multiple neutral DMIs that cannot, individually, cause speciation—a mechanism we call emergent speciation. We investigate emergent speciation using models of haploid holey adaptive landscapes—neutral networks—with recombination. We find that certain combinations of multiple neutral DMIs can lead to speciation. Furthermore, emergent speciation is a robust mechanism that can occur in the presence of migration, and of deviations from the assumptions of the neutral network model. Strong recombination and complex interactions between the DMI loci facilitate emergent speciation. These conditions are likely to occur in nature. We conclude that the interaction between DMIs may cause speciation.

**Author Summary:** Most species are kept distinct by incompatibilities between the genes they carry. These genetic incompatibilities cause hybrids between the species to have low fitness. Here we propose that combinations of several incompatibilities can collectively cause the origin of species, although they cannot do so acting alone—a mechanism we call emergent speciation. We use flat fitness landscapes with many holes in them to extend the classic Dobzhansky–Muller model, and capture the essence of the phenomenon. We find that emergent speciation can, indeed, occur through the combined effects of multiple genetic incompatibilities. Furthermore, the conditions that facilitate emergent speciation are likely to occur in nature. We conclude that the interaction between genetic incompatibilities may be a root cause of the origin of species.

## Introduction

Unravelling the ways in which reproductive barriers between populations arise and are maintained remains a central challenge of evolutionary biology. The Dobzhansky–Muller model posits that speciation is driven by intrinsic postzygotic reproductive isolation caused by incompatibilities between alleles at different loci [1–3]. The kinds of strong negative epistatic interactions envisioned by this model are common between amino acid substitutions within proteins [4, 5]. Furthermore, Dobzhansky–Muller incompatibilities (hereafter DMIs) have been shown to cause inviability or sterility in hybrids between closely related species, although the extent to which any particular DMI has actually caused speciation remains an open question [6–9].

In Fig. 1A, we illustrate a simple version of the evolutionary scenario originally proposed by Dobzhansky [1] with an incompatibility between neutral alleles at two loci (A and B) in a haploid—that is, a *neutral* DMI [10]. An ancestral population is fixed for the *ab* genotype. This population splits into two geographically isolated (allopatric) populations. One population fixes the neutral allele *A* at the A locus, whereas the other fixes the neutral allele *B* at the B locus. The derived alleles are incompatible: individuals carrying one of the derived alleles are fit but individuals carrying both of them are not. Upon secondary contact between the populations, this neutral DMI creates postzygotic isolation between the two populations: if *r* is the recombination rate between the loci, then *r/*2 of haploid F_1_ hybrids between individuals from the two populations are unfit (inviable or sterile).

**Figure 1.**
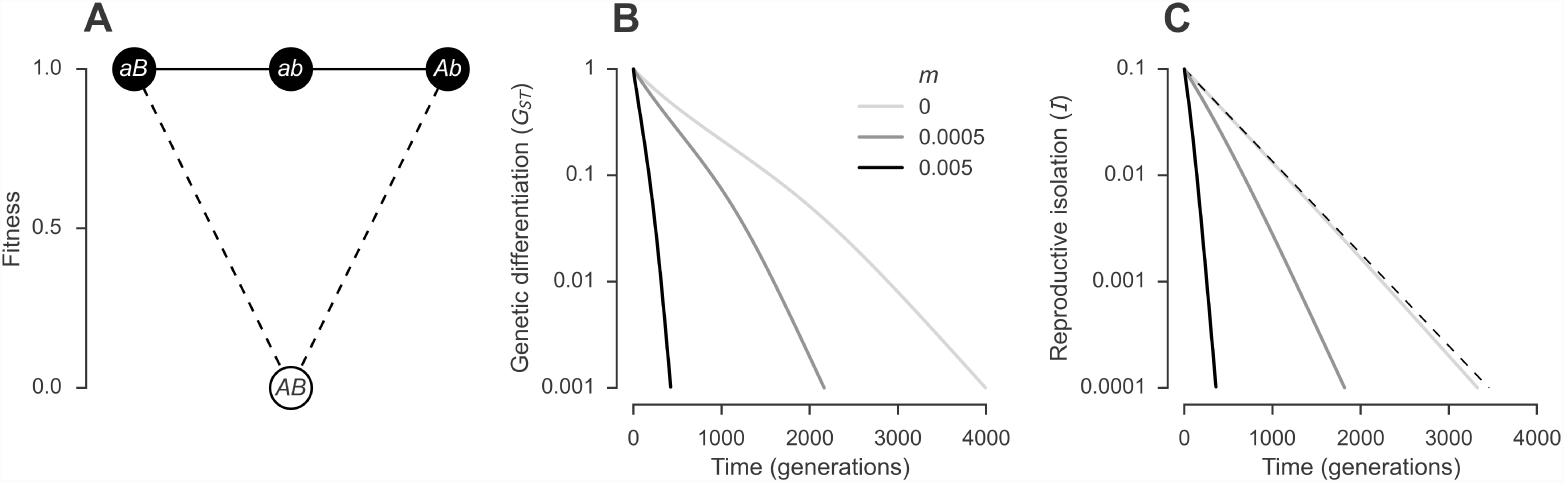
A neutral DMI between two loci is not sufficient to cause speciation. (A) Three haploid genotypes (closed circles) are viable and have equal fitness, and one genotype is inviable (open circle). The *K* = 3 viable genotypes form a neutral network. (B–C) The reproductive isolation generated by a neutral DMI between two diallelic loci does not persist in the face of either mutation or gene flow. Two populations start fixed for the *Ab* and *aB* genotypes, respectively, and are allowed to evolve with a mutation rate of *u* = 10^−3^ per locus per generation, a recombination rate of *r* = 0.2 between loci, and different levels of gene flow (*m*, proportion of a population consisting of migrants from the other population, each generation). Genotypes are allowed to mutate at both loci in a single generation. Initially, the genetic differentiation between the two populations is *G*_*ST*_ = 1 (B) and the degree of reproductive isolation is *I*_0_ = *r/*2 = 0.1 (C). Both *G*_*ST*_ and *I* decline over time. The dashed line in (C) shows *I*_*t*_ = *I*_0_ · *e*^−2*ut*^.

The neutral DMI described in the previous paragraph is unlikely to be an effective mechanism of speciation because it assumes that the populations diverge in perfect allopatry, and that the derived alleles go to fixation before secondary contact takes place. However, either mutation or gene flow can disrupt this process [10–12] (Fig. 1B and C): they lead to the production of individuals with the ancestral genotype (*ab*) and these individuals have an advantage because they are completely compatible with individuals carrying derived alleles (*Ab* and *aB*).

It is known that the reproductive barriers created by neutral DMIs can be strengthened in at least two ways. First, if selection favors the derived alleles—that is, if the DMI is *not* neutral [10, 13 –15]. This could happen if the derived alleles are involved in adaptation to different environments, a scenario known as ecological speciation [16, 17]. Second, if the two populations are prezygotically isolated. For example, the low fitness of hybrids can select against hybridization and cause the evolution of assortative mating between individuals carrying the same derived allele—a mechanism known as reinforcement [1, 18 –20]. Here we consider a new mechanism we call *emergent speciation*—that speciation emerges through the collective effects of multiple neutral DMIs that cannot, individually, cause speciation.

The majority of theoretical work on DMIs has relied on either population genetic models [10, 14, 15, 21–24], or models of divergence between populations [3, 25 –30]. Both classes of models include simplifying assumptions: the former consider only DMIs involving two or three loci, whereas the latter ignore polymorphism at the DMI loci. Both simplifications are problematic: reproductive isolation is often caused by multiple DMIs involving multiple loci [31–36], and many populations contain alleles involved in DMIs segregating within them [37–41]. Several studies that have attempted to overcome these simplifications have excluded DMIs [42], or have not represented DMIs explicitly [11, 43 –45], or have not considered neutral DMIs [46, 47] and, therefore, could not capture emergent speciation. Schumer et al. [48] considered multiple neutral DMIs between pairs of loci and found that they were ineffectual at driving hybrid speciation. We investigate emergent speciation using a haploid holey adaptive landscape model [12, 49]: the neutral network model [50, 51] with recombination [52, 53], which allows us to represent DMIs involving multiple loci, and to take into account genetic variation at those loci.

A neutral network [50, 51] is a network of viable genotypes connected by mutational accessibility. Two genotypes are mutationally accessible if one genotype can be obtained from the other through a single mutation. For example, Fig. 1A shows a neutral network where *aB* is connected to *ab* but not to *Ab*. All genotypes in the network are viable and have equal fitness. All genotypes outside the network are inviable but some may be mutationally accessible from genotypes in the network. For example, in the neutral network shown in Fig. 1A, *AB* is inviable, and it is accessible from both *aB* and *Ab*, but not *ab*.

Neutral networks extend the neutral DMI model to multiple loci [12, 49, 54]; a neutral network of *K* genotypes with *L* loci, each with *α* alleles can be constructed by taking the entire space of *α*^*L*^ genotypes and “removing” the *α*^*L*^ − *K* genotypes that carry incompatible combinations of alleles (e.g., the *A* and *B* alleles in the neutral network in Fig. 1A). The alleles of genotypes in the neutral network can be, for example, nucleotides, amino acids, insertions/deletions, or presence/absence of functional genes. Therefore, a neutral network can also be used to represent DMI-like scenarios such as reciprocal translocations [55, 56] and the degeneration of duplicate genes [25, 26, 57].

We show that neutral networks defined by multiple neutral DMIs can lead to the establishment of stable reproductive barriers between populations. Thus, emergent speciation can occur in principle. We also identify the two principal causes of emergent speciation: recombination and the pattern of interactions between DMI loci.

## Results

### A neutral DMI between two loci is not sufficient to cause speciation

Consider the neutral DMI illustrated in Fig. 1A. Initially, two allopatric populations are fixed for the *aB* and *Ab* genotypes, respectively. The populations are maximally genetically differentiated at the two loci (*G*_*ST*_ = 1). The degree of reproductive isolation between the two populations is *I* = *r/*2, the mean fitness of haploid F_1_ hybrids between individuals from the two populations (see Methods for definitions of both *G*_*ST*_ and *I*).

How stable is the reproductive barrier between the two populations? To address this question we begin by investigating the effect of mutation within populations. If the alleles at each locus can mutate into each other (*A↔a* and *B ↔b*) at a rate *u* per locus per generation, then the degree of reproductive isolation will decline exponentially (Fig. 1B and C, *m* = 0). Any amount of gene flow between the two populations will further accelerate the erosion of the reproductive barrier (Fig. 1B and C, *m >* 0).

The evolution of a stable reproductive barrier between two populations—that is, speciation—requires the existence of more than one stable equilibrium [24, 58]. A single neutral DMI between two diallelic loci is not sufficient to cause speciation because, in the presence of mutation (0 *< u <* 0.5), it only contains one stable equilibrium for any level of recombination [10, 12, 13] (S1 Text), and populations will gradually evolve toward this equilibrium (S1 Fig). Changes to the adaptive landscape can cause the appearance of two stable equilibria [10, 13]. For example, if the derived alleles confer an advantage (fitness: *w*_*aB*_ = *w*_*Ab*_ = 1 and *w*_*ab*_ = 1 − *s*), and if both *r* and *s* ≫ *u*, the genotype network will have two stable equilibria, with 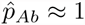 and 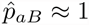 respectively (S2 Fig). Two populations in different equilibria will show a degree of reproductive isolation of: *I ≈ r*(1 + *s*)*/*2. Next we ask whether speciation can emerge through the collective effects of multiple neutral DMIs that cannot, individually, cause speciation.

### Speciation can emerge through the collective effects of multiple neutral DMIs that cannot, individually, cause speciation

To investigate if any neutral networks contain multiple stable equilibria we generated ensembles of 500 random neutral networks of *K* genotypes with *L* fitness loci and *α* alleles per locus for a range of values of *K*, *L* and *α*. To construct a random neutral network, we generated *K* random genotypes with *L* loci and one of *α* alleles at each locus (i.e., a “Russian roulette” model [12] with probability that a genotype is viable 𝒫 = *K/α^L^*), and kept the resulting network if it was connected. We ignored disconnected networks because, although they may contain multiple stable equilibria, shifts from one equilibrium to another require the simultaneous occurrence of multiple mutations and are, therefore, unlikely [12].

For each neutral network, we constructed populations with different initial genotype frequencies and allowed each population to evolve independently until it reached equilibrium. We then evaluated the stability of the resulting equilibria (see Methods). An exhaustive survey of all possible neutral networks defined on *L* = 3 diallelic loci (*α* = 2) revealed none containing multiple stable equilibria. However, some neutral networks with *L* = 4 diallelic loci, and with *L* = 3 triallelic loci (*α* = 3) contain multiple stable equilibria (Fig. 2A; S3 Fig).

**Figure 2.**
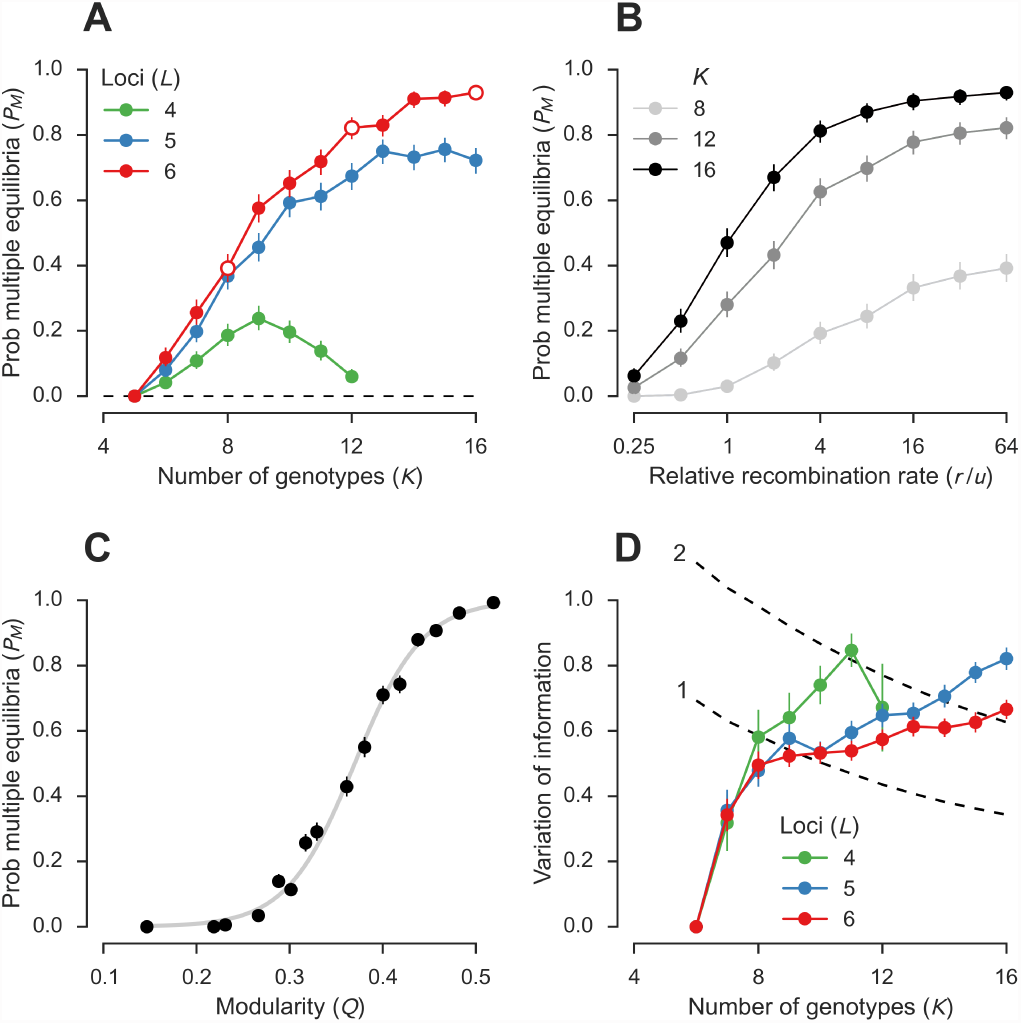
Speciation can emerge through the collective effects of multiple neutral DMIs that cannot, individually, cause speciation. (A) Probabilities that neutral networks contain multiple stable equilibria (*P*_*M*_) based on ensembles of 500 random connected neutral networks for each combination of numbers of genotypes (*K*) and diallelic loci (*L*). Error bars are 95% confidence intervals (modified Jeffreys intervals [59]). The presence of multiple equilibria was evaluated under a mutation rate of *u* = 10^−3^ per locus per generation (up to two mutations were allowed per genotype per generation) and a recombination rate of *r* = 0.064 between adjacent loci (up to *L* − 1 crossovers were allowed between two genotypes per generation); relative strength of recombination, *r/u* = 64. (B) The existence of multiple stable equilibria depends on the relative strength of recombination, *r/u*. The three neutral network ensembles denoted by open circles in (A) were reanalysed for a range of recombination rates while keeping the mutation rate constant (*u* = 10^−3^). See S4 Fig for the effect of changing *u* while keeping *r/u* constant. (C) Modular neutral networks are more likely to show multiple stable equilibria (see Table 1). Values are estimates and 95% CIs of 16 bins of 10^3^ random neutral networks based on pooling the data used in (A). The line shows a logistic regression model (the line encompasses a 95% confidence region based on 10^4^ bootstrap samples). (D) Modules inferred by maximizing *Q* [see (C)] show good agreement with modules inferred from the multiple stable equilibria of neutral networks. Dashed lines show the expected variation of information associated with changing the module membership of one or two genotypes.

Populations evolving independently to different stable equilibria become genetically differentiated (Fig. 3A) and partially reproductively isolated (Fig. 3C) from each other. In networks with *K* = 16 genotypes with *L* = 6 loci, the *average* level of reproductive isolation achieved between different stable equilibria is *I* = 15%, that is, over half of its maximum possible value (Fig. 3C). Thus, speciation can emerge through the collective effects of multiple neutral DMIs that cannot, individually, cause speciation. Next we investigate the mechanistic basis of emergent speciation.

**Figure 3.**
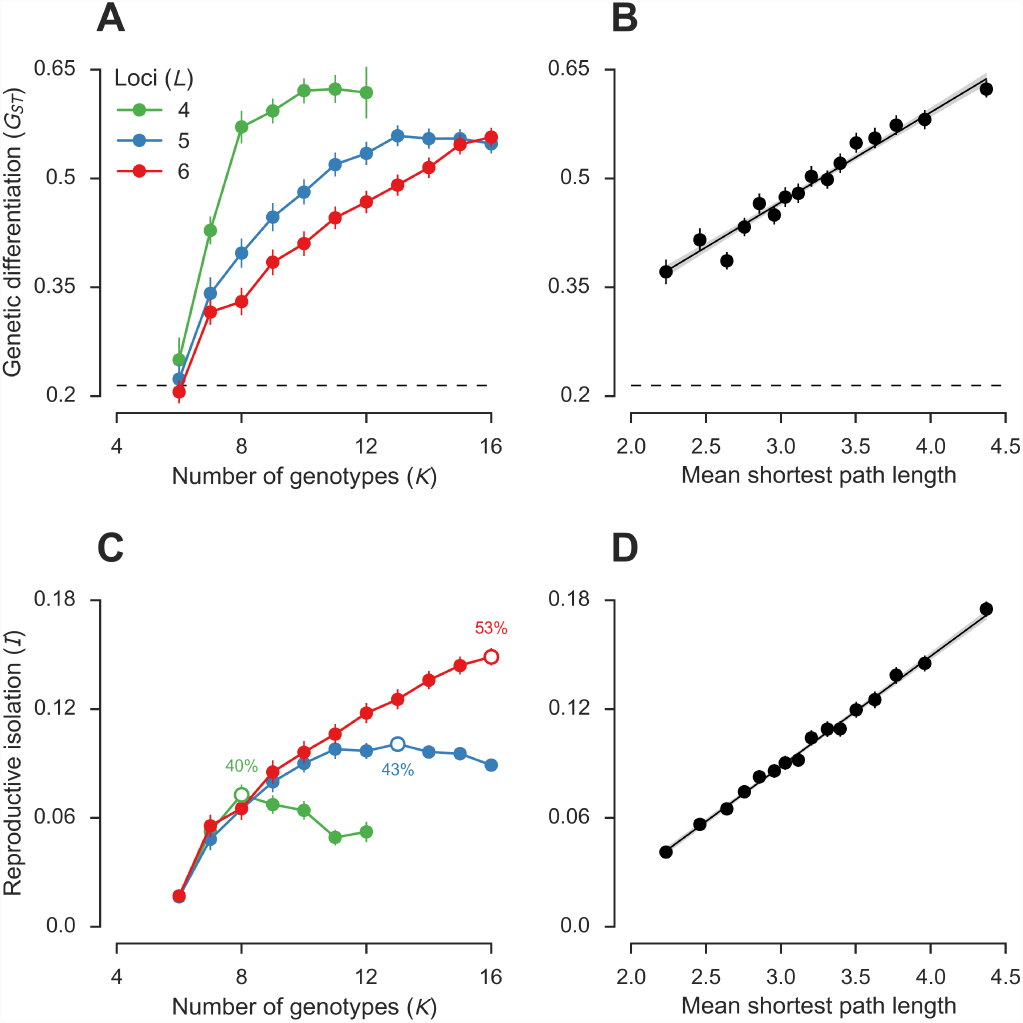
Emergent speciation can lead to high levels of genetic differentiation and reproductive isolation. Genetic differentiation (*G*_*ST*_, A–B), and degree of reproductive isolation (*I*, C–D) between populations at different equilibria. If a neutral network contained more than two stable equilibria, the maximum pairwise *I* and corresponding *G*_*ST*_ were used. The equilibria were calculated under the population genetic parameters described for Fig. 2A (*r/u* = 64). (A, B) Dashed line indicates the average *G*_*ST*_ observed in 905 species of animals and plants [61, 62]. (A, C) Values are means and 95% CIs of *G*_*ST*_ and *I* based on the 7013 neutral networks containing two or more stable equilibria in the ensembles in Fig. 2A. Open circles in (C) indicate the ensembles of random neutral networks with a certain number of loci (*L*) showing the highest average value of *I*. The numbers show the average proportion of the maximum possible value of *I* achieved in each ensemble. (B, D) Both *G*_*ST*_ and *I* are positively correlated with mean shortest path length (see Table 1). Values are estimates and 95% CIs of 16 bins of *∼* 400 random neutral networks each based on pooling the data used in (A) and (B). The lines show regression models; the gray regions indicate 95% confidence regions based on 10^4^ bootstrap samples.

### Emergent speciation requires that recombination be strong relative to mutation

In the absence of recombination between the loci defining a neutral network, there is only one stable equilibrium [51]. The genotype frequencies at equilibrium are given by the leading eigenvector of the mutation matrix **M** [see Methods, Eq. (3)] [51]. With recombination, however, multiple stable equilibria can occur (Fig. 2A). Thus, emergent speciation requires recombination.

To quantitatively investigate the relationship between the existence of multiple stable equilibria and the recombination rate between fitness loci (*r*), we calculated the probability, *P*_*M*_, that a random neutral network from an ensemble shows multiple stable equilibria for a range of values of *r* and *u*. We found that *P*_*M*_ increases with the relative strength of recombination *r/u*, and that the rate of increase differs among ensembles (Fig. 2B). Changing both *r* and *u* while keeping *r/u* constant has only a small effect on Explanatory power (%) of linear models with each property (see Methods) as independent variable and an indicator of emergent speciation as dependent variable. “Other” refers to properties of the genotypes that constitute the neutral network, not just of the network itself. The probability that a neutral network contains multiple stable equilibria (*P*_*M*_) was modelled by logistic regression using the data on 1.6 × 10^4^ random neutral networks summarized in Fig. 2A; the explanatory power of a network property was measured by the coefficient of discrimination [60]. The genetic differentiation (*G*_*ST*_) and degree of reproductive isolation (*I*) among equilibria were modelled by linear regression on the 7,013 random neutral networks containing two or more stable equilibria summarized in Fig. 3A and C, respectively; the explanatory power of a network property was measured by the coefficient of determination (*R*^2^). A sign in the “Direction” column indicates that the property is significantly correlated (*P <* 0.001) with all indicators of emergent speciation in the same direction. Values in bold, highlight the network property with the greatest explanatory power for a given indicator of emergent speciation.

*P*_*M*_ (S4 Fig).

The effect of the relative strength of recombination on emergent speciation can be seen more clearly for a specific neutral network. Fig. 4A shows one of the smallest neutral networks showing multiple stable equilibria (one of the random neutral networks in the *K* = 6, *L* = 3 and *α* = 3 ensemble summarized in S3 Fig). It is defined by a lethal allele (*B*_3_) and 8 pairwise DMIs: *A*_2_–*B*_2_ (i.e., *A*_2_ and *B*_2_ are incompatible), *A*_2_–*C*_1_, *A*_2_–*C*_3_, *A*_3_–*B*_2_, *A*_3_–*C*_1_, *A*_3_–*C*_3_, *B*_1_–*C*_1_ and *B*_1_–*C*_3_; it has a mean DMI order of *ω* = 1.57 [see Methods, Eq. (2)]. The equilibria of this neutral network follow a supercritical pitchfork bifurcation [63] with the relative strength of recombination, *r/u*, as control parameter (Fig. 4B; S5 Fig B). When recombination is weak relative to mutation (*r/u <* 2) the neutral network contains only one stable equilibrium regardless of initial conditions. Above a critical relative strength of recombination (*r/u >* 2) there are two stable equilibria and one unstable equilibrium and populations evolve to the different equilibria depending on initial conditions (Fig. 4B and C; S2 Text). The stable equilibria are defined by high frequency of the *B*_1_ (blue) and *B*_2_ (red) allele, respectively (Fig. 4B and C; S2 Text).

**Figure 4.**
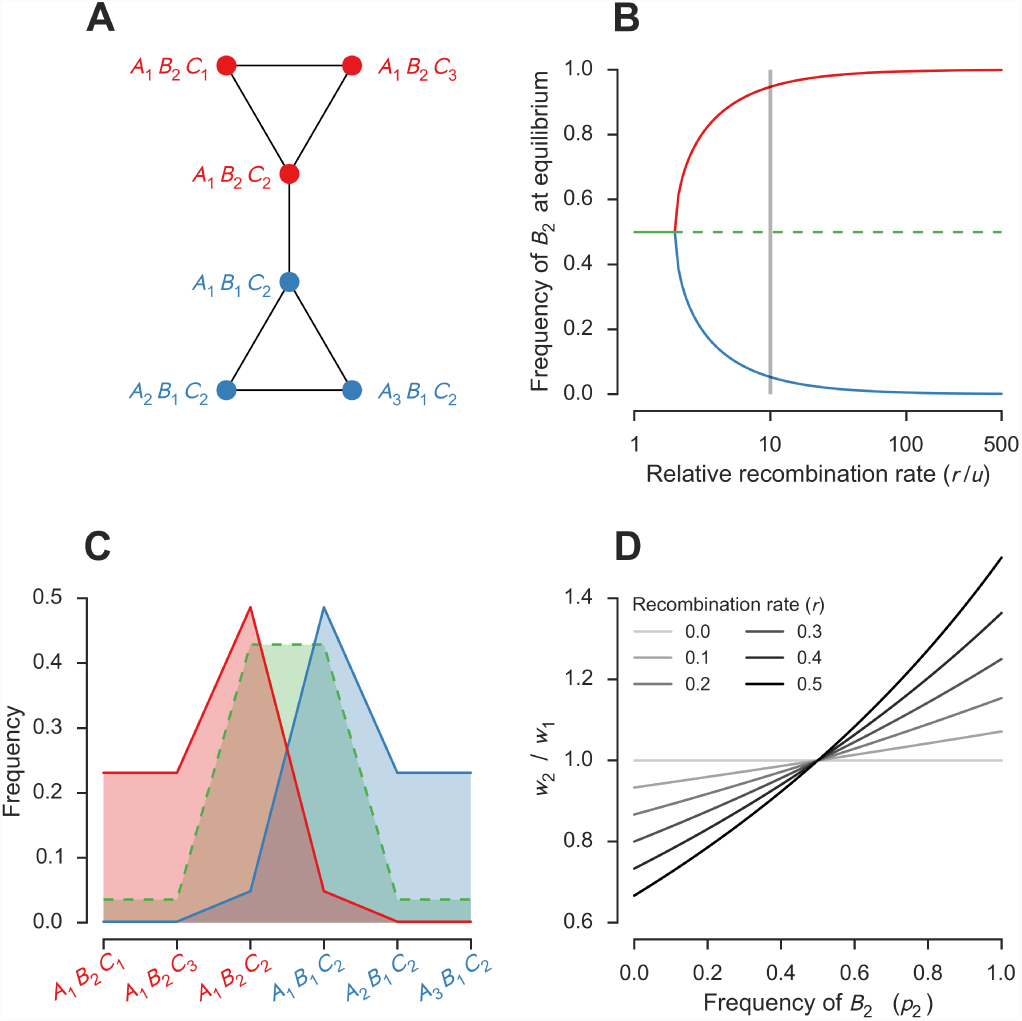
Example of a neutral network supporting emergent speciation. (A) Neutral network of the *K* = 6 genotypes with *L* = 3 loci and *α* = 3 alleles per locus containing either the *A*_1_*B*_2_ haplotype (red) or the *B*_1_*C*_2_ haplotype (blue). The colors relate to the equilibria as explained below. (B) The frequency of the *B*_2_ allele at equilibrium follows a supercritical pitchfork bifurcation [63] with the relative strength of recombination, *r/u*, as control parameter: when *r/u <* 2.0, there is a single stable equilibrium with the *B*_2_ allele at a frequency of 50% (solid green line); when *r/u >* 2.0, there are two stable equilibria, with high and low frequencies of *B*_2_, respectively (red and blue lines), and an unstable equilibrium with *B*_2_ at a frequency of 50% (dashed green line). The figure shows *u* = 10^−3^. See S5 Fig B for the effect of *u* on the critical value of *r/u*. (C) Genotype frequencies at equilibrium for *r/u* = 10 [gray line in (B)]. Populations initially fixed for any of three genotypes shown in red (i.e., containing *B*_2_) evolve to the stable equilibrium in red, whereas populations initially fixed one of the 3 genotypes shown in blue (i.e., containing *B*_1_) evolve to the stable equilibrium in blue. Populations showing equal frequencies of all genotypes evolve to the unstable equilibrium in green. (D) The two stable equilibria are maintained in part by positive frequency-dependent selection. Lines show the selective advantage of the *B*_2_ allele (red) relative to the *B*_1_ allele (blue) in populations with different frequencies of *B*_2_ [Eq. (1)]. The *B*_2_ allele is favored when it is in the majority, and its relative marginal fitness (*w*_2_*/w*_1_) increases with its frequency (*p*_2_); the strength of frequency-dependent selection increases with *r*. Unless otherwise stated, we used the same population genetic parameters as in Fig. 2A.

The gradual increase of *P*_*M*_ with *r/u* in random neutral networks (Fig. 2B) indicates that the relative strength of recombination required for emergent speciation to occur differs among neutral networks within an ensemble. Next we ask which network properties, if any, promote emergent speciation?

### Modular neutral networks are more likely to lead to emergent speciation

Comparison of *P*_*M*_ among the ensembles of random neutral networks shows that *P*_*M*_ increases with network size, *K* (Fig. 2A; S3 Fig; Table 1). We have never found a random connected neutral network with *K ≤* 5 genotypes with multiple stable equilibria (*P*_*M*_ *≈* 0), regardless of the values of *L* and *α*. In contrast, networks with *K* = 16 genotypes defined by *L* = 6 diallelic loci, show *P*_*M*_ *≈* 93% (Fig. 2A). *P*_*M*_ also increases with *L* and *α*, that is, with the size of the genotype space from which the neutral network is sampled (Fig. 2A; S3 Fig; Table 1).

**Table 1.** Many properties of neutral networks correlate with emergent speciation.

| Property | Direction | $P_M$ | $G_{ST}$ | $I$ |
| --- | --- | --- | --- | --- |
| *Network* |  |  |  |  |
| Algebraic connectivity | — | 39.8* | 4.5* | 19.9* |
| Coefficient of variation in degree |  | < 0.1 | 4.1* | 1.8* |
| Connectance | — | 47.1* | 13.1* | 24.5* |
| Degree assortativity | — | 0.1* | 2.7* | 0.2* |
| Estrada index | + | 11.1* | 9.6* | 6.4* |
| Mean degree |  | 0.3* | 4.0* | < 0.1 |
| Mean shortest path length | + | 48.8* | **18.1\*** | **38.6\*** |
| Modularity ($Q$) | + | **54.9\*** | 9.5* | 30.3* |
| Network size ($K$) | + | 28.9* | 13.0* | 15.7* |
| Number of modules | + | 19.0* | 5.7* | 8.6* |
| Spectral radius |  | < 0.1 | 1.9* | < 0.1 |
| Square clustering coefficient |  | 6.5* | 0.5* | < 0.1 |
| *Other* |  |  |  |  |
| Mean DMI order ($\omega$) | + | 10.2* | 19.1* | 1.6* |
| Mean Hamming distance | + | 46.7* | 11.5* | 36.5* |
| Number of loci ($L$) | | 12.3* | 3.3* | 9.5* |

Although *P*_*M*_ is correlated with many properties of the neutral networks, the best predictor we have found of whether a neutral network shows multiple stable equilibria is the modularity of the network (Fig. 2C; Table 1). A module is a group of densely interconnected genotypes showing relatively few connections to genotypes outside the module [64]. Modules inferred by considering only the structure of a neutral network are good predictors of the equilibria—that is, species—found in the neutral network (Fig. 2D). For example, the neutral network shown in Fig. 4A contains two modules shown in blue and red, which correspond exactly to the two stable equilibria observed when recombination is strong relative to mutation.

Modularity contributes to emergent speciation in two ways. First, modularity generates a mutational bias towards staying in a module, as opposed to leaving it. For example, consider the *B*_2_ (red) module in the neutral network shown in Fig. 4A: *A*_1_*B*_2_*C*_1_ and *A*_1_*B*_2_*C*_3_ can only generate *B*_2_ (red) mutants, and *A*_1_*B*_2_*C*_2_ generates twice as many *B*_2_ (red) mutants as *B*_1_ (blue) mutants. Introducing a mutational bias towards leaving the *B*_2_ module opposes emergent speciation (S5 Fig A). The mutational biases associated with modularity are not, however, sufficient to cause emergent speciation in the absence of recombination, which brings us to the second way in which modularity promotes emergent speciation. Suppose that most of the population occupies a particular module of the neutral network. Although mutations can generate individuals outside the dominant module, recombination between those “outsiders” and individuals from the dominant module will produce a disproportionate number of offspring inside the dominant module. This creates positive frequency-dependent selection for genotypes in that module. For example, in the neutral network shown in Fig. 4A, if each *B* allele is evenly distributed between all viable genotypes containing it then the selective advantage of the *B*_2_ allele relative to the *B*_1_ allele is given by (ignoring the effect of mutation):

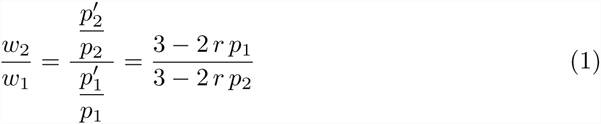

where *w*_*i*_ and *p*_*i*_ are the marginal fitness and frequency of the *B*_*i*_ allele, respectively. Eq. (1) is illustrated in Fig. 4D. The *B*_2_ allele is favored relative to *B*_1_ (*w*_2_*/w*_1_ *>* 1) when it is in the majority (*p*_2_ *>* 0.5, Fig. 4D). The strength of this frequency-dependent selection is proportional to the recombination rate (Fig. 4D).

Modularity is also strongly correlated with other indicators of emergent speciation: genetic differentiation (*G*_*ST*_) and reproductive isolation (*I*) (Table 1). However, the best predictor we have found of both *G*_*ST*_ and *I* is the mean shortest path length between genotypes (Fig. 3B and 3D). In other words, emergent speciation is associated with a neutral network “syndrome” characterized by (Table 1): large size, sparse connectivity (low connectance, low algebraic connectivity), looseness (high mean shortest path length, high Estrada index), and modularity (high *Q*, high number of modules). However, the topology of a network is *not* sufficient to determine *P*_*M*_: the precise pattern of linkage between loci also influences whether a particular neutral network shows multiple stable equilibria (S7 Fig). Next we investigate the robustness of emergent speciation to deviations from the assumptions of the neutral network model.

### Emergent speciation can occur in the presence of gene flow

If two allopatric populations evolve independently to the different stable equilibria of the neutral network in Fig. 4A, they will become genetically differentiated and reproductively isolated to an extent that also depends on *r* (Fig. 4B and D; S6 Fig A and B).

The reproductive barrier created by the neutral network in Fig. 4A can persist in the presence of gene flow (S6 Fig). Introducing gene flow weakens the degree of genetic differentiation and of reproductive isolation at equilibrium, and increases the critical value of *r/u* required for the persistence of a reproductive barrier. However, the maximum migration rate between two populations that allows the reproductive barrier to persist is low, of the order of the mutation rate (S6 Fig C). Stable differentiation can occur in a stepping-stone model [65] with higher local migration rates (S9 Fig D), but the resulting reproductive barrier does not slow down the spread of a neutral allele at an unlinked locus appreciably [11] (Table 2).

**Table 2.** Multiple neutral DMIs can generate strong barriers to gene flow.

| Scenario | $T_{25}$ | $b$ |
| --- | --- | --- |
| DMI with selection (S2 Fig A) |  |  |
| $s = 0.05$ | 1,556 | 1.027 |
| $s = 0.1$ | 1,583 | 1.045 |
| $s = 0.25$ | 1,632 | 1.077 |
| $s = 0.5$ | 1,690 | 1.116 |
| $s = 0.75$ | 1,742 | 1.150 |
| $s = 1$ | 1,790 | 1.182 |
| Neutral network |  |  |
| Fig. 4A | 1,513 | 0.999 |
| S8 Fig A | 1,778 | 1.174 |

Larger neutral networks, however, can generate much stronger reproductive barriers, capable of withstanding substantial gene flow. For example, the neutral network shown in S8 Fig A contains three stable equilibria (S8 Fig D). This was one of the random neutral networks in the *K* = 11 and *L* = 5 ensemble summarized in Fig. 2A. Populations at the equilibria at opposite ends of the network can show high levels of genetic differentiation and reproductive isolation. If the fitness loci are unlinked (*r* = 0.5), then 50% of F_1_ hybrids between two populations at equilibrium are inviable, and the maximum migration rate between two populations that allows the reproductive barrier to persist is almost two orders of magnitude higher than the mutation rate (*m ≈* 0.0943). In a stepping-stone model, this neutral network can slow down the spread of a neutral allele at an unlinked locus to a greater extent than a single DMI with strong selection for the derived alleles (S9 Fig C and E; Table 2). Thus, emergent speciation can, in principle, occur in either allopatry or parapatry. Next we investigate the extent to which emergent speciation depends on the assumptions of the model we have been using.

### Emergent speciation is a robust mechanism

The neutral network model analysed so far includes two central assumptions: neutrality within the network, and complete inviability outside it. To investigate the extent to which emergent speciation depends on these assumptions we have relaxed them in turn for the neutral network shown in Fig. 4A. First, emergent speciation is robust to some variation in fitness among the genotypes in a neutral network. If there is free recombination between the fitness loci then there will be two stable equilibria even if the fitness of one of the four outer genotypes is doubled relative to that of the other genotypes in the network (S5 Fig C). Second, provided the disadvantage of leaving the neutral network is substantial, partial DMIs still allow the existence of stable—albeit weaker—reproductive barriers (S5 Fig D).

We have also assumed infinite population size in all our analyses. But emergent speciation can also occur in finite populations. If there is free recombination between the fitness loci, populations of moderate size (*N ≈* 1000) are expected to remain in equilibrium for thousands of generations (Fig. 5). We conclude that emergent speciation is a robust mechanism. Next we investigate the genetic basis of emergent speciation.

**Figure 5.**
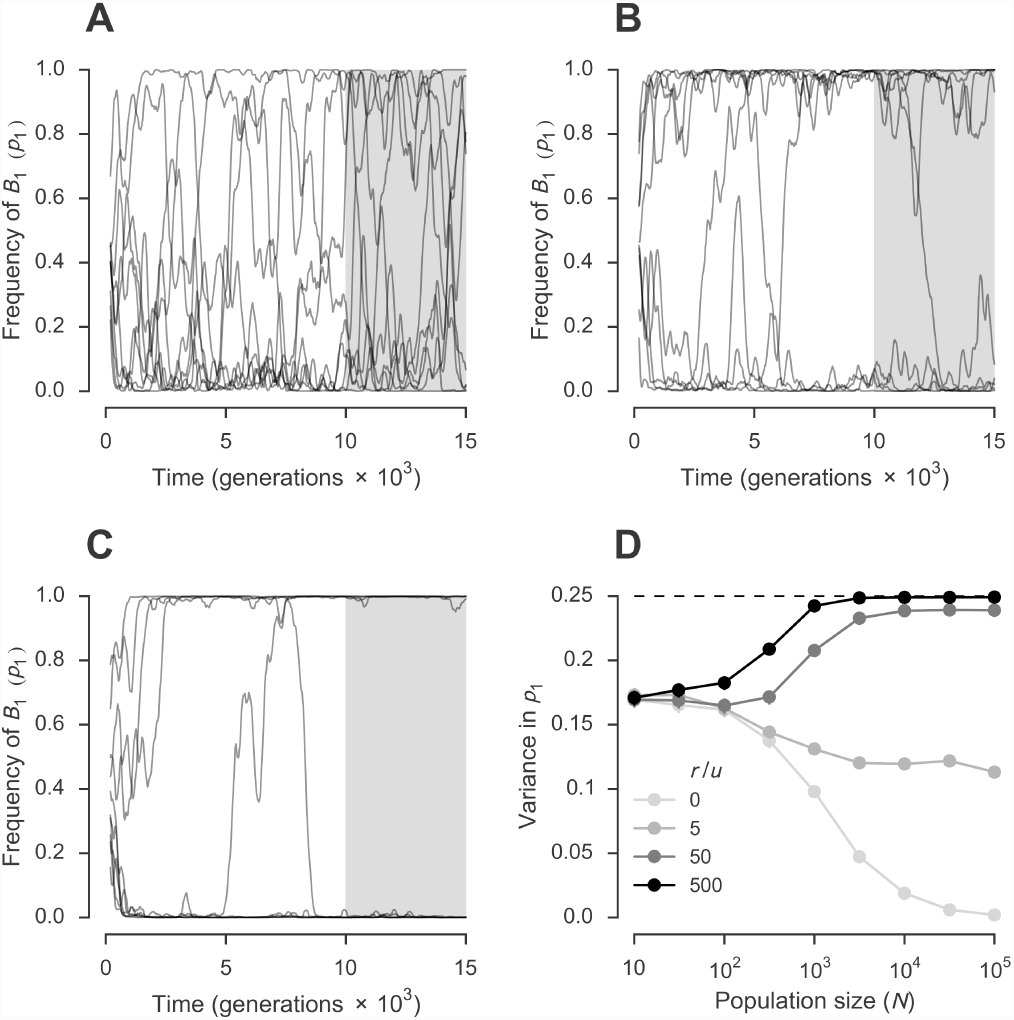
Emergent speciation can occur in finite populations. So far, all analyses have assumed infinite population size. (A–C) Ten populations of *N* = 10^3^ individuals were allowed to evolve on the neutral network shown in Fig. 4A under a range of relative strength of recombination: *r/u* = 5 (A), 50 (B), and 500 (C). Values show moving averages of the frequency of the *B*_1_ allele, *p*_1_, with a window of 200 generations. Initially, each population was sampled with replacement from a population with equal numbers of each genotype. (D) Variance in *p*_1_ among populations. For each population, we calculated the average *p*_1_ between generation 10^4^ and 1.5 × 10^4^ [gray sectors in (A–C)]. Values are variances in *p*_1_ based on 10^3^ populations. Error bars are 95% CIs based on 10^4^ bootstrap samples. The dashed line shows the maximum possible value of the variance in *p*_1_. Unless otherwise stated, we used the same population genetic parameters as in Fig. 2A.

### Complex DMIs promote emergent speciation

We refer to the number of loci involved in a DMI as its *order*. DMIs of order two are designated *simple*, whereas DMIs of order three or greater are designated *complex* [3, 30, 66]. A neutral network of a certain size defined by a certain number of loci can be specified by either a few low-order DMIs or many high-order DMIs (Fig. 6). This is because a low-order DMI causes the inviability of more genotypes than a higher-order one.

**Figure 6.**
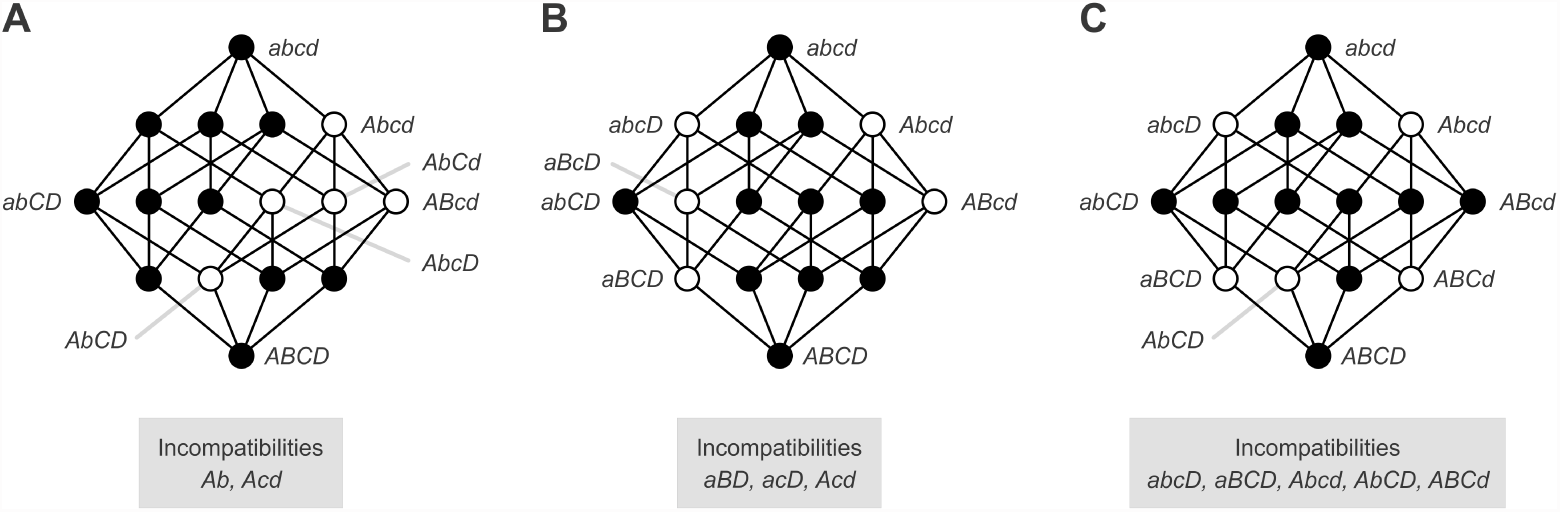
A neutral network of a certain size defined by a certain number of loci can be specified by either a few low-order DMIs or many high-order DMIs. Three neutral networks containing *K* = 11 genotypes with *L* = 4 diallelic loci. Neutral network determined by one simple DMI and one complex DMI of order 3 [mean DMI order, *ω* = 2.2, see Methods, Eq. (2)]. (B) Neutral network determined by three complex DMIs of order 3 (*ω* = 3). This is the experimentally inferred neutral network of all the mutational intermediates between the 5S ribosomal RNA sequences of *Vibrio proteolyticus* (lowercase alleles) and *V. alginolyticus* (uppercase alleles) [67]. (C) Neutral network determined by five complex DMIs of order 4 (*ω* = 4).

In random networks, there is a positive correlation between mean DMI order (*ω*) and indicators of emergent speciation, specially genetic differentiation (Table 1). To investigate this relationship more closely, we chose the ensemble with the greatest variance in *ω* in Fig. 2A (*K* = 12 and *L* = 5) and generated new ensembles of random neutral networks over the full range of possible values of *ω*. To avoid sampling biases, all the networks considered had different network topologies. These ensembles of random networks confirm the strong positive relationship between mean DMI order and the probability that a network contains multiple stable equilibria. Thus, higher order DMIs promote emergent speciation (Fig. 7).

**Figure 7.**
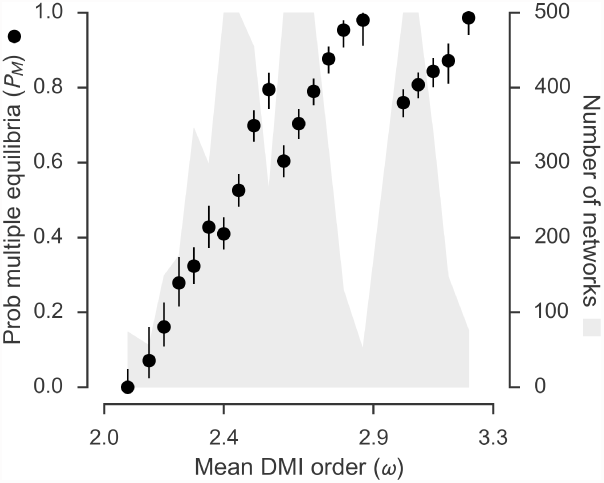
Higher-order DMIs promote emergent speciation. Neutral networks determined by higher-order DMIs are more likely to show multiple stable equilibria. Probabilities that neutral networks of *K* = 12 genotypes with *L* = 5 diallelic loci contain multiple stable equilibria (*P*_*M*_) based on ensembles of up to 500 random connected neutral networks with particular mean DMI order, *ω*. Each ensemble contains neutral networks with different topologies. Neutral networks defined by lethal alleles (i.e., effectively involving *L <* 5 loci) were excluded from the analysis. Ensembles containing fewer than 500 networks include one network for each existing topology (based on an exhaustive search of all networks with *K* = 12 and *L* = 5). Ensembles with fewer than 50 networks were pooled with neighboring ensembles until the resulting ensemble had at least 50 networks. Error bars are 95% CIs. Population genetic parameters are the same as in Fig. 2A.

## Discussion

Our main result is that, when it comes to multiple neutral DMIs, the whole can be greater than the sum of its parts. Although a single neutral DMI cannot lead to the evolution of stable reproductive isolation, the collective effects of combinations of multiple neutral DMIs can lead to the evolution of strong barriers to gene flow between populations—a mechanism we call emergent speciation.

### Causes of emergent speciation

Emergent speciation has two interacting causes: recombination and the pattern of interactions between DMI loci. Increasing the relative strength of recombination between DMI loci promotes emergent speciation in at least three ways. First, it causes the appearance of multiple equilibria (Fig. 2B; Fig. 4B; S2 Text). Recombination had been shown to generate multistability in other evolutionary models [68–74], although earlier studies of the evolutionary consequences of recombination in neutral networks did not detect multiple equilibria [52, 53]. Second, it increases genetic differentiation between populations at the different equilibria (S6 Fig A). This pattern is consistent with the observation that increasing *r* reduces variation within a population at equilibrium in a neutral network [52, 53, 75]. Third, it increases the degree of reproductive isolation between populations at different equilibria (S6 Fig B). This is because, in our model, recombination can generate inviable hybrids and consequently is the predominant source of selection. High *r* between fitness loci has been shown to promote speciation in other models [10, 18].

Recombination may also *oppose* emergent speciation by reducing the probability of a stochastic shift between stable equilibria [12, 23, 58] (Fig. 5). Therefore, the probability of emergent speciation may be maximized at intermediate strengths of recombination.

Not all patterns of interaction between DMI loci can produce emergent speciation regardless of the strength of recombination. Multiple DMIs involving only pairs of loci (i.e., simple DMIs) rarely lead to emergent speciation. For example, we found 9 neutral networks with different topologies with *K* = 12 and *L* = 5 specified entirely by simple DMIs (*ω* = 2), none of which had multiple stable equilibria (Fig. 7; but note that the neutral network in Fig. 4A is specified entirely by DMIs among pairs of loci). A recent study reported consistent findings in a model of hybrid speciation in a diploid [48]. More complex interactions between the DMI loci, however, promote emergent speciation (Table 1; Fig. 7). For example, *∼* 76% of neutral networks with *K* = 12 and *L* = 5 specified by DMIs of order *ω* = 3 on average had multiple stable equilibria (Fig. 7). Complex DMIs have only rarely been modeled in previous studies [3, 30], which could explain why emergent speciation had not been discovered before.

### Conditions for the existence of emergent speciation

The causes of emergent speciation outlined in the previous section suggest three conditions for the existence of emergent speciation in nature: (i) multiple DMIs should segregate within populations, (ii) the loci involved in these DMIs should not all be tightly linked, and (iii) at least some of these DMIs should have high order.

There is strong support for all three conditions. Many populations contain alleles involved in DMIs segregating within them [37–41]. Recently, Corbett-Detig and colleagues [39] found evidence that multiple simple DMIs involving unlinked loci are currently segregating within natural populations of *Drosophila melanogaster*. They surveyed a large panel of recombinant inbred lines (RILs) and found 22 incompatible pairs of alleles at unlinked loci in the RILs; of the 44 alleles, 27 were shared by two or more RILs, indicating that multiple DMIs are polymorphic within natural populations.

Corbett-Detig et al. did not attempt to identify higher-order DMIs, presumably because they lacked the statistical power to do so [39]. Although high-order epistasis is thought to be widespread, it is more difficult to detect it experimentally because it requires the construction and phenotypic analysis of more genotypes [30, 54, 76, 77]. Nevertheless, complex DMIs have been discovered in introgression studies [30]. For example, Orr and Irving [78] investigated the sterility of male F_1_ hybrids between the USA and Bogota subspecies of *D. pseudoobscura* and found that it is caused by an incompatibility between loci in both chromosomes 2 and 3 of USA and loci in at least three different regions of the X chromosome of Bogota—a DMI of order *ω ≥* 5 (of which only two loci might be tightly linked).

The principal conditions for emergent speciation are met in natural populations. The details of the model we used imply two additional conditions. We consider them in the next section.

### The “holeyness” of adaptive landscapes

The neutral network model we used makes two extreme assumptions about the fitness of genotypes: neutrality within the network and complete inviability outside it. These assumptions are not met universally. “In-network” neutrality will occur if every DMI is neutral. However, many DMI loci appear to have experienced positive selection during their evolutionary history [6–8]. Similarly, many DMIs are only mildly deleterious, rather than lethal [31, 36, 39]. Despite this, emergent speciation will operate in nature either if the fitness assumptions are frequently met in nature, or if emergent speciation is robust to departures from these assumptions. We shall consider each possibility in turn.

Beyond anecdotal evidence, we do not know how often either assumption is met in nature. Having said that, both assumptions are realistic. A substantial fraction of new nonsynonymous mutations is nearly neutral (i.e., have *N*_*e*_ *|s| <* 1, where *N*_*e*_ is the effective population size and *s* is the fitness effect of the mutation): *∼* 25%, *∼* 6% and 19% for *Saccharomyces cerevisiae* [79], *D. melanogaster* [80, 81] and *Mus musculus castaneus* [81], respectively. If even a small proportion of these mutations can participate in DMIs, that would imply the existence of large numbers of neutral DMIs. Likewise, DMIs often cause approximately complete sterility or inviability. For example, Presgraves [31] screened *∼* 70% of the *D. simulans* autosomal genome for regions that reduce male viability when they are hemyzogous in the presence of a hemyzogous *D. melanogaster* X chromosome and discovered 20 approximately lethal DMIs.

Setting aside whether the fitness assumptions are met in nature, our results suggest that emergent speciation is robust to deviations from the assumptions. Emergent speciation does not require lethal DMIs. As long as the the disadvantage of “falling” off the neutral network is substantial, partial DMIs are still expected to lead to the evolution of stable reproductive barriers (S5 Fig D). Emergent speciation is also robust to some variation in fitness among the genotypes in a neutral network (S5 Fig C). Therefore, in-network neutrality can approximate more complex scenarios where selection is weak or variable over time and/or space, or effective population sizes are small [12]. We propose that speciation should still be classed as emergent if the alleles involved in the DMIs are not under sustained positive selection.

### Emergent speciation in diploids

In diploids, a single neutral DMI is also not sufficient to cause speciation [10]. Therefore, there may be a diploid equivalent of emergent speciation. Strictly, our model is only valid for haploids. However, extrapolating from the analysis of a single DMI between two loci by Bank, Bürger and Hermisson [10], we expect that our results will apply broadly to diploids provided that the DMIs are recessive, such that selection acts primarily against recombinant F_2_ hybrids. Presgraves estimates that there are *∼* 8× more recessive than dominant lethal DMIs between *D. melanogaster* and *D. simulans* [31].

If DMIs are dominant or codominant, selection will act primarily against F_1_ hybrids, a mechanism that is not captured in our model. Thus, if dominant or codominant neutral DMIs can cause their own version of emergent speciation, it is likely to have different causes from the emergent speciation described in our study; for example, it may require weak, rather than strong, recombination [10].

### Predictions from emergent speciation

Our results suggest four predictions. If emergent speciation is operating in nature, we predict that DMIs fixed between species should: (i) be less tightly linked and (ii) have higher order (*ω*), on average, than DMIs segregating within species. These predictions cannot be tested at present because we do not have unbiased estimates of either the number of DMIs or their order in divergence and polymorphism in any system. Some known DMIs indicate that these predictions could be met in nature. For example, Davis et al. showed that loci in both chromosomes 2 and 3 of *D. mauritiana* are incompatible with loci in the X chromosome of *D. simulans*, causing female sterility [82]—a DMI of order *ω ≥* 3 involving unlinked loci.

Prediction (ii) has been made independently by others based on a different argument: that complex DMIs evolve more easily than simpler DMIs because they allow a greater proportion of the possible evolutionary paths between the common ancestor and the evolved genotypes containing the incompatibility [3, 66]. Fraïsse et al. tested this mechanism using simulations and concluded that it is unlikely to be effective [30].

One complication for experimental tests of predictions (i) and (ii) is that, just because there is a DMI fixed between two species, it does not mean that it was involved in speciation, emergent or otherwise; it could simply be a by-product of divergence after speciation has occurred by other means [6–9]. At present, we do not know how to distinguish between causal and noncausal DMIs. Therefore, testing these hypotheses is likely to be most effective if speciation has occurred recently.

The precise pattern of recombination—that is, linkage—between loci can also promote emergent speciation (S7 Fig). This result indicates that certain genomic rearrangements may facilitate emergent speciation. We predict (iii) that if emergent speciation is operating in nature, genomic rearrangements will correlate with speciation. Chromosomal speciation models make similar predictions [83, 84]. Unlike those models, however, our model does not require that the genomic rearrangements affect the fitnesss of hybrids or suppress recombination [83, 84].

Finally, we predict (iv) that two kinds of genomic rearrangements—translocations and gene duplications—participate in emergent speciation more directly. Below, we focus on gene duplication but essentially the same argument can be made about reciprocal translocations [55, 56]. Gene duplications followed by reciprocal degeneration or loss of duplicate copies in different lineages can act just like DMIs [25, 26], despite not involving an epistatic interaction [57]. Gene duplications, degenerations and losses are common and a substantial fraction of gene degenerations and losses are likely to be effectively neutral [57, 85 –87]. If the duplicates are essential, then genotypes carrying insufficient functional copies will be completely inviable. For example, Bikard et al. [37] showed that this mechanism underlies a simple DMI between two strains of *Arabidopsis thaliana*. Following whole genome duplications, multiple gene degenerations or losses occur [57, 85, 86, 88], and the duplicates tend to be unlinked. Both reciprocal translocations and reciprocal degeneration or loss of duplicate genes appear to have contributed to the diversification of yeasts [55, 56, 88].

### Speciation and evolution on holey adaptive landscapes

Gavrilets [12, 49] introduced the idea that speciation is a consequence of evolution on holey adaptive landscapes (e.g., neutral networks) containing ridges of viable genotypes connecting distant genotypes [89]. He imagined populations moving through these ridges and speciation occurring when two populations evolve to genetic states separated by holes [12].

Our results broadly support Gavrilets’ vision, but suggest two ways in which it can be refined. First, speciation is not equally likely to occur on every holey adaptive landscape. The *structure* of the network of genotypes over which populations evolve also plays an important role: emergent speciation is promoted large size, sparse connectivity, looseness, and modularity of the genotype network. Species are defined by modules in the genotype network. Second, recombination is critical. Gavrilets’ work on holey adaptive landscapes [12, 49] has tended to downplay the importance of recombination. (As, indeed, has most earlier work on neutral networks [51, 90 –92].) Our results suggest that both the structure of the genotype network and the strength of recombination determine whether speciation will occur.

Our results have broader evolutionary implications. The neutral network model has played an important role in the study of the evolution of robustness and evolvability [51, 90 –93]. Our finding that recombination promotes the appearance of multiple stable equilibria in neutral networks has clear implications for the evolution of robustness and evolvability that deserve further investigation. For example, Wagner [94] has argued that recombination helps explore genotype space because it causes greater genotypic change than mutation. However, our results suggest that, depending on the structure of the neutral network, large sexual populations can get trapped in stable equilibria, therefore restricting their ability to explore genotype space (evolvability).

## Conclusion

We have discovered a new mechanism of speciation: that it emerges from the collective effects of multiple neutral DMIs that cannot, individually, cause speciation. The conditions that promote emergent speciation are likely to occur in nature. We conclude that the interaction between DMIs may be a root cause of the origin of species. Continued efforts to detect DMIs [32, 33, 36] and to reconstruct real neutral networks [67] (Fig. 6B) will be crucial to establishing the reality and importance of emergent speciation.

## Methods

### Neutral networks

Organisms are haploid and carry *L* loci with effects on fitness. Each locus can have one of *α* alleles. Out of the possible *α*^*L*^ genotypes, *K* are viable, with equal fitness, and the remaining (*α*^*L*^ - *K*) genotypes are completely inviable. The *K* genotypes define a neutral network, where genotypes are connected if one genotype can be obtained from the other through a single mutation (i.e., they differ at a single locus).

### Statistics

#### Algebraic connectivity

Second smallest eigenvalue of the Laplacian matrix of the network [95]. Calculated using NetworkX [96].

#### Connectance

Total number of connections between genotypes as a proportion of the maximum possible number of connections between genotypes, *K*(*K -* 1).

#### Degree assortativity

A measure of the correlation of the degree of linked genotypes [95]. Calculated using NetworkX [96].

#### Estrada index

A centrality measure [97]. Calculated using NetworkX [96].

#### Mean and coefficient of variation of degree

Mean and coefficient of variation (standard deviation divided by the mean) of the degree distribution, respectively [95]. The degree of a genotype is the number of its viable mutational neighbors. Calculated using NetworkX [96].

#### Mean DMI order

Let *x*_*i|*𝒩_ be the number of DMIs of order *i ≥* 2 (i.e., involving a particular combination of alleles at *i* loci) in a neutral network, 𝒩; *x*_*i|*𝒩_ can also be defined for *i* = 1, in which case it measures the number of lethal alleles in 𝒩. (For simplicity, we shall refer to *i* as DMI order.) Let 𝒩_*i*_ be the neutral network implied by the *x*_*i|*𝒩_ DMIs of order *i*. If the networks 𝒩_*i*_ and 𝒩 are the same, then network *N* only involves DMIs of order *i* or fewer. Let *X*_*i|*𝒩_ = *x*_*L|*𝒩_*i* be the number of DMIs of order of the number of loci, *L*, in 𝒩_*i*_. The mean DMI order, *ω*, of neutral network *N* is given by:

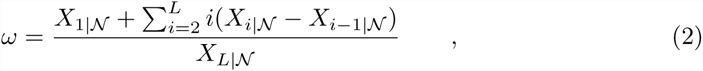

where *X*_*L|*𝒩_ = *x*_*L|*𝒩_ = *α*^*L*^ - *K*, the number of inviable genotypes in the neutral network 𝒩 (note that 𝒩_*L*_ = 𝒩).

#### Mean Hamming distance

Mean number of loci at which pairs of genotypes carry different alleles. Genotypes connected in the neutral network are at a Hamming distance of 1.

#### Mean shortest path length

Mean number of steps along the shortest path between pairs of genotypes [95]. Calculated using NetworkX [96].

#### Modularity and number of modules

Modularity (*Q*) is a measure of the extent to which the network displays modules [64]. Calculated using igraph [98] based on an exhaustive search for the partition that maximizes *Q* among all possible partitions of the network.

#### Network size

Number of genotypes (*K*).

#### Spectral radius

Leading eigenvalue of adjacency matrix [95]. The mean degree of a population at equilibrium in the absence of recombination if genotypes are only allowed to mutate at one locus per generation [51]. Calculated using NumPy [99].

#### Square clustering coefficient

The probability that two viable mutational neighbors of a genotype *g* share a common viable mutational neighbor other than *g*. Calculated using NetworkX [96].

### Evolution

Evolution on a neutral network was modeled by considering an infinite-sized population of haploid organisms reproducing sexually in discrete generations. The state of the population is given by a vector of frequencies 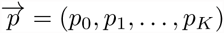, where *p*_*i*_ is the frequency of genotype *i*. Genotypes outside the network are ignored because they are completely inviable [51]. Individuals mate at random with respect to genotype to form a transient diploid that undergoes meiosis to produce haploid descendants. Selection takes place during the haploid phase. Mating, recombination, mutation and selection cause the population to evolve according to the equation:

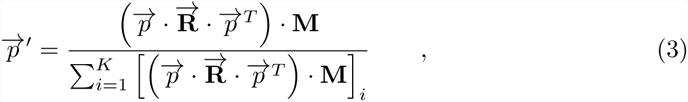

where 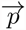 and 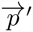 are the states of the population at generations *t* and *t* + 1, respectively.

**M** is the mutation matrix such that entry *M*_*ij*_ is the mutation rate from genotype *i* to genotype *j* per generation. The diagonal elements of **M** (*M*_*ii*_) represent the probability that genotype *i* does not mutate (including to inviable genotypes outside the neutral network). Values of *M*_*ij*_ are set by assuming that each locus mutates with probability *u* and that a genotype can only mutate simultaneously at up to two loci.

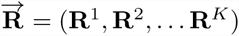 is a vector of recombination matrices such that entry 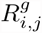 of matrix **R**^*g*^ is the probability that a mating between individuals of genotypes *i* and *j* generates an individual offspring of genotype *g* (more precisely, 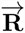 is a tensor of rank 3). Up to *L -* 1 crossover events can occur between two genotypes with probability 0 *≤ r ≤* 0.5 per interval. The recombination rate *r* is assumed to be the same for all pairs of adjacent loci. If *r* = 0.5, then there is free recombination between all loci.

### Equilibria

#### Identification

Given a neutral network, population genetic parameters *u* and *r*, and a set of initial genotype frequencies 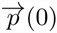, the population was allowed to evolve until the root-mean-square deviation of the genotype frequencies in consecutive generations was RMSD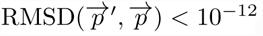. The final genotype frequencies were identified as an equilibrium 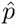

Multiple initial conditions (i.e., genotype frequencies) were tested: (i) fixed for each of the *K* genotypes in turn, (ii) even distribution over all *K* genotypes, (iii) equilibrium genotype frequencies in the absence of recombination (*r* = 0), and (iv) 10 independent sets of random frequencies sampled uniformly over the (*K -* 1)-simplex. Two equilibria, 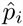 and 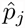, were judged identical if RMSD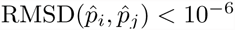. Only one of a set of identical equilibria was counted. This procedure does not guarantee the discovery of all equilibria.

#### Stability analysis

To evaluate the stability of an equilibrium 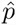, we initialized a population with frequencies 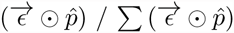, where 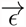 is a vector of random numbers uniformly distributed between 0.99 and 1.0, and ⨀ represents the Hadamard product of two vectors. We then allowed the population to evolve to equilibrium again, without further perturbation. If the new equilibrium was identical to the original equilibrium, it was classified as *stable*.

In a large subset of the equilibria studied (including all the equilibria found in networks with *K ≤* 12 showing multiple equilibria), stability was validated by calculating the eigenvalues of the Jacobian of 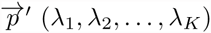 at 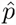; the equilibrium was classified as stable if *∀*_*i*_*|λ*_*i*_*| <* 1.

#### Modules

In addition to the modularity analysis described above (see “Network statistics”), modules were inferred from multiple stable equilibria in the following way: a genotype *g* was assigned to a module corresponding to a stable equilibrium 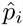 if the frequency of *g* was higher in 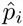 than in any other stable equilibrium. The resulting “equilibrium” modules were compared to the modules inferred by maximizing *Q* using the variation of information metric [100].

### Gene flow

#### Two populations

Gene flow was modeled as symmetric migration between two populations. Migration occurs at the beginning of each generation, such that a proportion *m* of each population is composed of immigrants from the other population. Then random mating, recombination and mutation take place within each population, as described above.

#### Stepping-stone model

A stepping-stone model [65] was used to measure the rate of spread of a neutral allele across a reproductive barrier [11]. A number *n* of populations are arranged in a line. Every generation a proportion 2*m* of a population emigrates to its two neighboring populations (except populations 1 and *n*, which have only one neighbor, so only *m* of each of them emigrate) (see S9 Fig A). Note that, unlike the stepping-stone model studied by Gavrilets [14], our implementation allows the genotype frequencies of terminal populations (1 and *n*) to vary. Therefore, all *n* populations may evolve to the same equilibrium. When *n* = 2 this model reduces to the gene flow model described in the previous section.

#### Genetic differentiation

*G*_*ST*_ = 1 - *H*_*S*_ /*H*_*T*_ was used to measure the genetic differentiation between two populations at a locus, where *H*_*S*_ is the average gene diversity of the two populations, and *H*_*T*_ is the gene diversity of a population constructed by pooling the two populations [21]. The gene diversity of a population at a locus is defined as 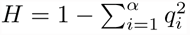, where *q*_*i*_ is the frequency of allele *i* and *α* is the number of alleles. Values of *G*_*ST*_ can vary between 0 (two populations with the same allele frequencies) and 1 (two populations fixed for different alleles). The overall genetic differentiation between two populations was quantified as the average *G*_*ST*_ over all loci. If all genotypes in the neutral network contain the same allele at a locus, that locus is excluded from the calculation of average *G*_*ST*_.

#### Reproductive isolation

The degree of reproductive isolation between two populations is defined as [58, 101]: 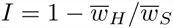, where 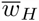 is the mean fitness of haploid F_1_ hybrid offspring from crosses between individuals from the two populations, and 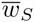 is the average of the mean fitnesses of the individual populations. The calculation of 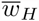 and 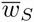 only takes into account the contribution of recombination, and ignores mutation. Values of *I* can vary between 0 (e.g., if all F_1_ hybrids are viable, or the populations are undifferentiated, or *r* = 0) and 1 (all F_1_ hybrids are inviable). More precisely, the maximum possible value of *I* is the total proportion of recombinants in a cross between two genotypes: 1− (1− *r*)^*L*−1^, where *L* is the number of fitness loci and *r* is the recombination rate between adjacent loci.

## Acknowledgments

We thank N. Barton, T. Cooper, J. Cuesta, J. Krug, A. Kalirad, S. Manrubia, I. Nemenman, and D. Weissman, for helpful discussions. A. Kalirad and I. Patanam contributed to coding, testing, and documentation.

## Supporting Information

**S1 Text**

**S1 Text A neutral DMI between two diallelic loci is not sufficient to cause speciation.** Analysis of the number of stable equilibria for the neutral network in Fig. 1A under two evolutionary scenarios: (a) no recombination, and (b) strong recombination between the fitness loci.

**S2 Text**

**Emergent speciation requires that recombination be strong relative to mutation.** Analysis of the number of stable equilibria for the neutral network in Fig. 4A under two evolutionary scenarios: (a) no recombination, and (b) strong recombination between the fitness loci.

## S1 Text: A neutral DMI between two diallelic loci is not sufficient to cause speciation

Consider the neutral network with *K* = 3 genotypes illustrated in Fig. 1A. There are two fitness loci (A and B) with two alleles each (*A/a* and *B/b*, respectively). *aB*, *ab* and *Ab* are viable (fitness, *w* = 1) and *AB* is inviable (*w* = 0).

We analyse the number of stable equilibria in this neutral network under two evolutionary scenarios: (a) no recombination, and (b) strong recombination between the fitness loci.

### (a) No recombination

We assume that the fitness loci are tightly linked (*r* = 0). The state of the population is defined by the vector of genotype frequencies: 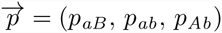.

If we assume that the mutation rate is *u* per locus per generation, and that only one locus is allowed to mutate per generation, we get the following mutation matrix:

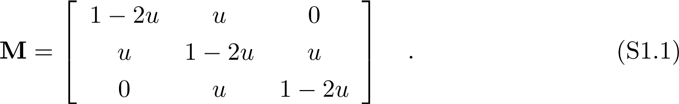

The genotype frequencies in the following generation 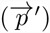 are given by:

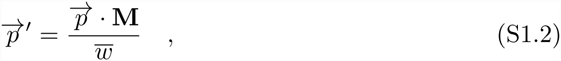

where 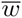 is the mean fitness of the population:

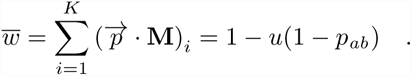

Expanding S1.2 we get:

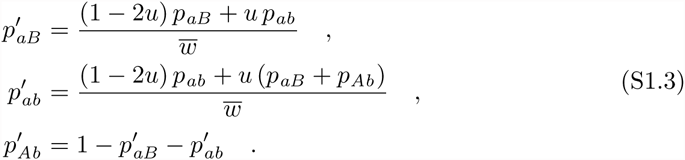

Solving the system of equations 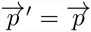 (S1.3) *we get one equilibrium:*

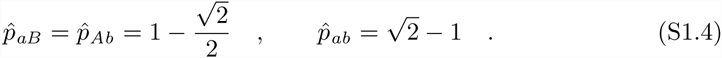

This equilibrium is the leading eigenvector of **M** (S1.1) [1].

The eigenvalues of the Jacobian at S1.4 are:

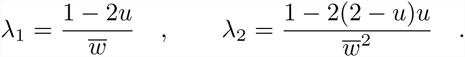

Equilibrium S1.4 is stable if *∀*_*i*_ *|λ*_*i*_*| <* 1. This condition is met if 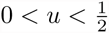.

### (b) Strong recombination

We assume that recombination between the fitness loci is strong enough that the population is always in linkage equilibrium [2]: *p*_*ij*_ = *p*_*i*_ *× p*_*j*_, where *i* is an allele at the A locus and *j* is an allele at the B locus. Thus, the state of the population is defined by the vector of *allele* frequencies: 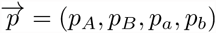

If we assume that the mutation rate is *u* per locus per generation, the allele frequencies in the following generation 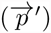 are given by:

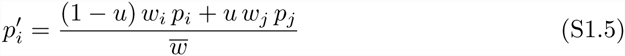

where *i* and *j* are both alleles at the same locus, *p*_*i*_ is the frequency of allele *i*, *w*_*i*_ is the marginal fitness of allele *i*:

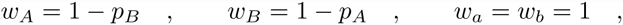

and 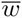 is the mean fitness of the population:

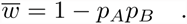

Solving the system of equations 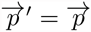 (S1.5), we find one equilibrium:

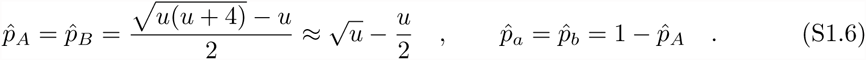

The genotype frequencies at equilibrium are:

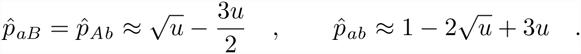

The eigenvalues of the Jacobian at S1.6 are:

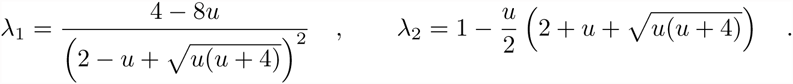

Equilibrium S1.6 is stable if *∀*_*i*_ *|λ*_*i*_*| <* 1. This condition is met if 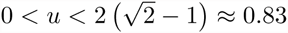.

### (c) Conclusion

A single neutral DMI between two diallelic loci contains only one equilibrium under both evolutionary scenarios, regardless of the mutation rate (0 *< u <* 0.5). Therefore, it cannot cause speciation. These results are consistent with those reported by Rutschman [3], Gavrilets [4] (pp. 125–8), and Bank et al. [5]. (Only Gavrilets considered mutation; Bank et al. used a continuous-time model.)

## S2 Text: Emergent speciation requires that recombination be strong relative to mutation

Consider the neutral network with *K* = 6 genotypes illustrated in Fig. 4A. There are three fitness loci (A, B and C) with three alleles each (*A*_1_*/A*_2_*/A*_3_, *B*_1_*/B*_2_*/B*_3_ and *C*_1_*/C*_2_*/C*_3_, respectively). Genotypes containing either the *A*_1_*B*_2_ haplotype or the *B*_1_*C*_2_ haplotype are viable (*w* = 1) and all other genotypes are inviable (*w* = 0).

### (a) No recombination

We assume that the DMI loci are tightly linked (*r* = 0). The state of the population is defined by the vector of genotype frequencies, 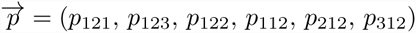, where *p*_*ijk*_ refers to the frequency of the *A*_*i*_*B*_*j*_*C*_*k*_ genotype.

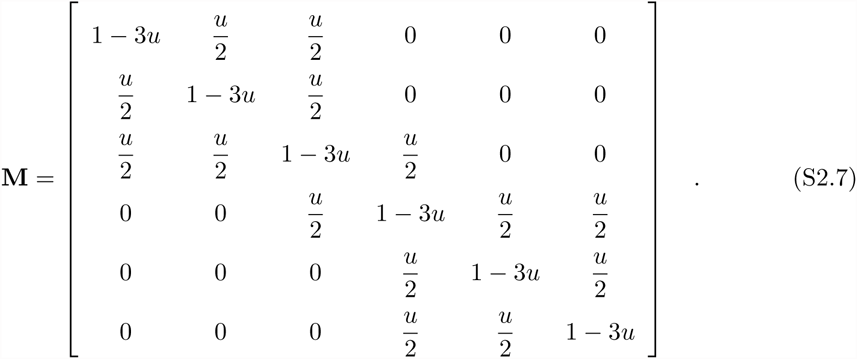

The genotype frequencies in the following generation are given by Equation S1.2:

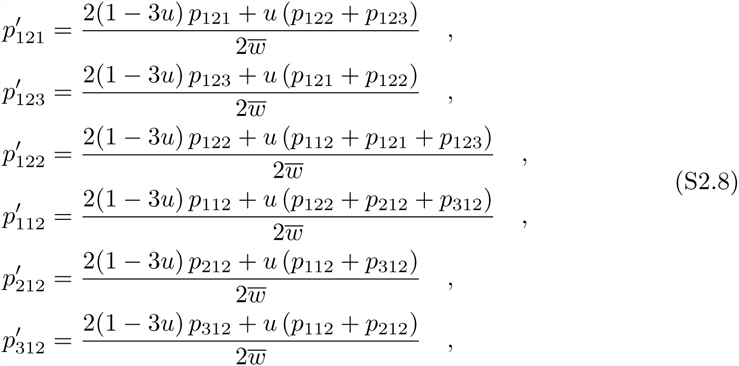

where 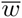 is the mean fitness of the population:

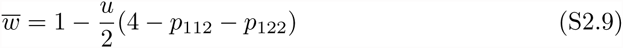

Solving the system of equations 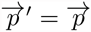 (Equations S2.8) we find one equilibrium:

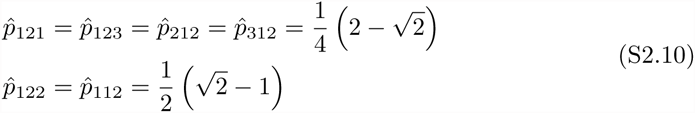

This equilibrium is the leading eigenvector of **M** (S2.7) [1].

The eigenvalues of the Jacobian at S2.10 are:

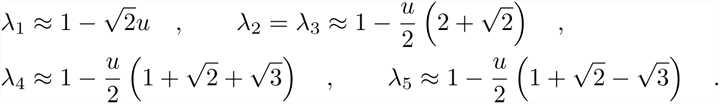

(First order Taylor expansions around *u* = 0.)

Equilibrium S1.6 is stable if *∀*_*i*_ *|λ*_*i*_*| <* 1. This condition is met if 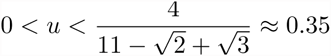 (obtained from the exact values of *λ*_*i*_, not the approximate ones).

### (b) Strong recombination

We assume that recombination between the fitness loci is strong enough that the population is always in linkage equilibrium [2]: *p*_*ijk*_ = *pA*_*i*_ *× pB*_*j*_ *× pC*_*k*_. Thus, the state of the population is defined by the vector of *allele* frequencies: 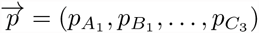. If we assume that the mutation rate is *u* per locus per generation, the allele frequencies in the following generation 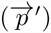 are given by:

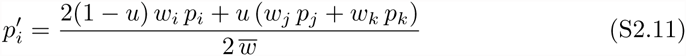

where *i*, *j* and *k* are alleles at the same locus, *p*_*i*_ is the frequency of allele *i*, *w*_*i*_ is the marginal fitness of allele *i*:

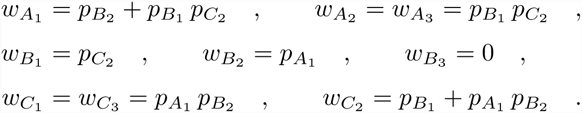

and 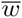 is the mean fitness of the population:

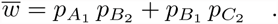

Solving the system of equations 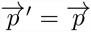 (S2.11), *we find three equilibria.*

**Equilibrium I** The allele frequencies at equilibrium are:

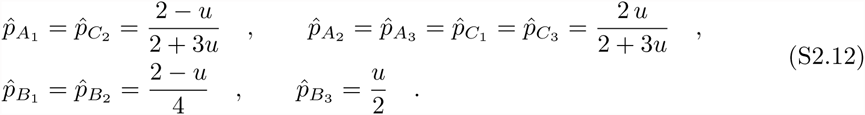

The genotype frequencies at equilibrium are:

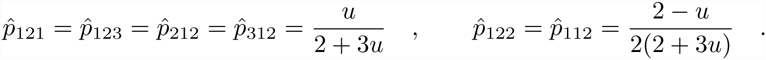

The eigenvalues of the Jacobian at S2.12 are:

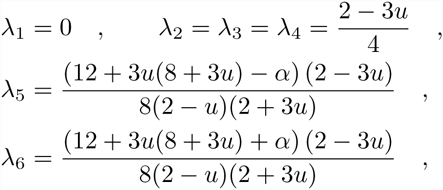

where 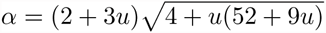.

Equilibrium S2.12 is stable if *∀*_*i*_ *|λ*_*i*_*| <* 1. This condition is met if 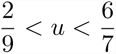.

**Equilibrium II** The allele frequencies at equilibrium are:

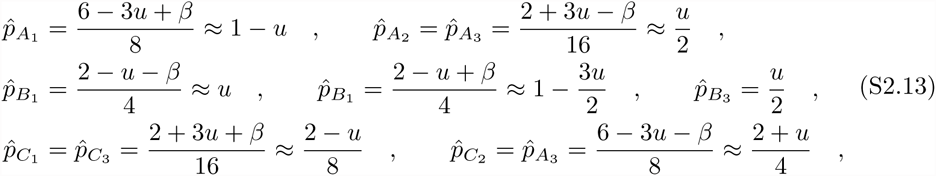

where 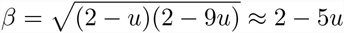.

The genotype frequencies at equilibrium are:

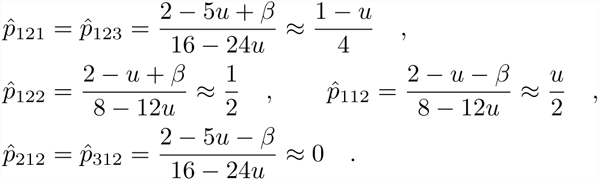

The eigenvalues of the Jacobian at S2.13 are:

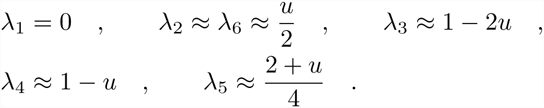

(First order Taylor expansions around *u* = 0.)

Equilibrium S2.13 is stable if *∀*_*i*_ *|λ*_*i*_*| <* 1. This condition is met if 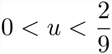 (obtained from the exact values of *λ*_*i*_, not the approximate ones).

**Equilibrium III** This equilibrium is symmetrical with equilibrium II according to the following pattern of correspondence between alleles:

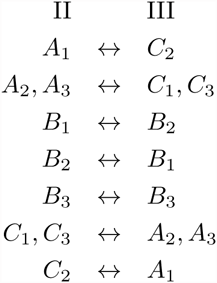

Thus, this equilibrium is also stable if 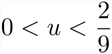.

**Mean fitness** At a given mutation rate, the mean fitness at equilibrium is highest for the stable equilibrium/equilibria:

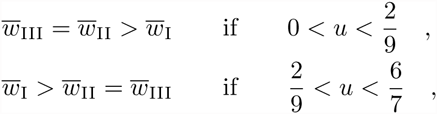

where 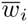 is the mean fitness at equilibrium *i*.

### (c) Conclusions

- In the absence of recombination between the fitness loci there is only one stable equilibrium regardless of the mutation rate (0 < *u* ≲ 0.35).
- When both recombination and mutation are strong 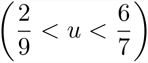 there is only one stable equilibrium. This equilibrium becomes unstable when mutation is weaker 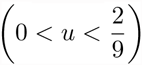.
- When recombination is strong and mutation is relatively weak 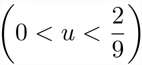 there are two stable equilibria and one unstable equilibrium (see previous point).
- Assuming that the strong recombination regime corresponds to *r ≈* 0.5, the critical relative strength of recombination for the existence of multiple stable equilibria is *r/u ≈* 2, in agreement with the numerical results on the full model shown in Fig. 4B and S5 Fig B.
- When recombination is strong populations are expected to evolve to a stable equilibrium where population mean fitness is maximized.

## Supporting Figures

**Figure S1.**
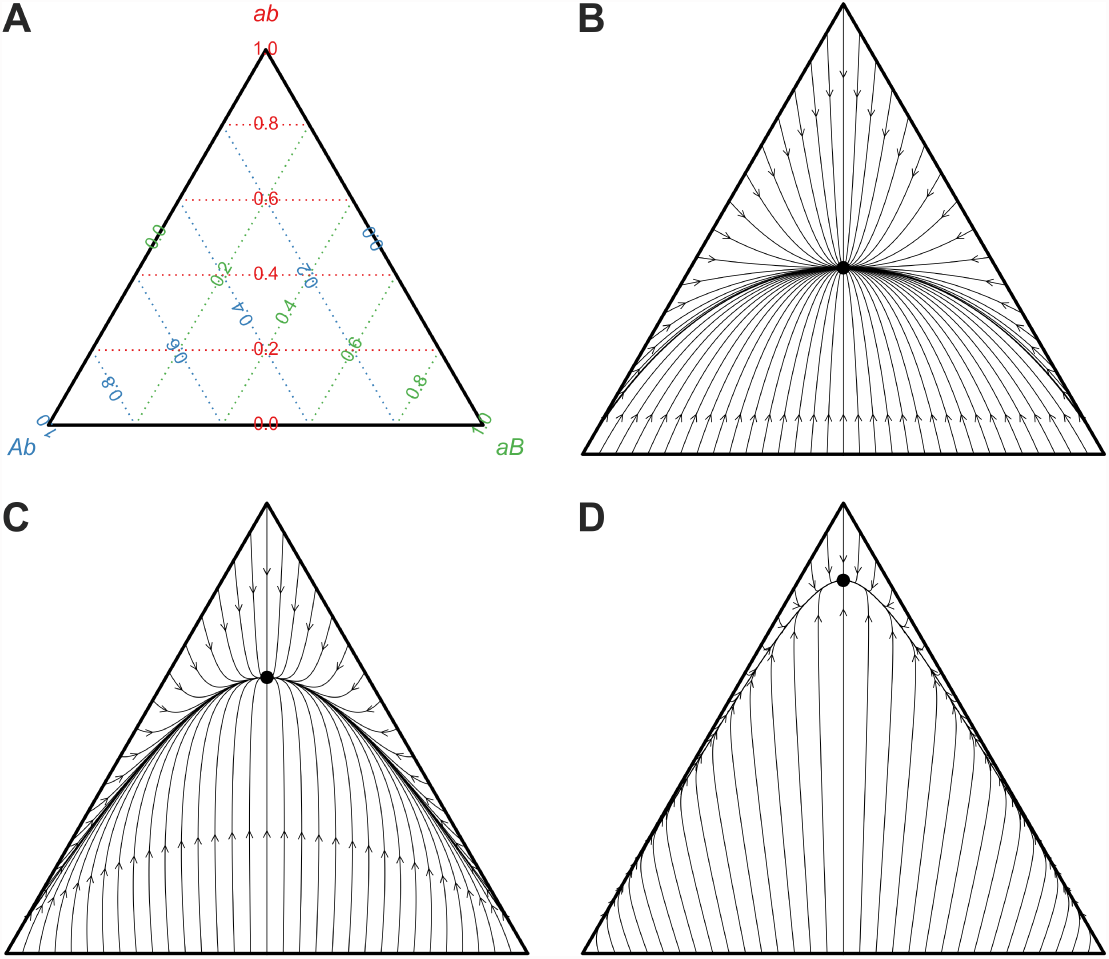
A neutral DMI between two diallelic loci contains only one stable equilibrium. (A) The frequencies of the three genotypes (*ab*, *Ab* and *aB*, see Fig. 1A) in a population are represented as a single point in a ternary plot. (B) Evolutionary trajectories without recombination (*r* = 0) of populations starting at initial frequencies such that at least one of the genotypes is absent (i.e., the edges of the triangle). All trajectories converge to a single stable equilibrium (solid circle) with frequencies 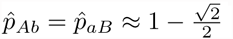.and 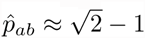 (S1 Text). Arrowheads mark the genotype frequencies after 100 generations of evolution. (C–D) Evolutionary trajectories with recombination rates of *r* = 0.01 and 0.1, respectively. As the recombination rate increases, the frequency of *ab* at equilibrium 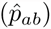 increases, and populations approach equilibrium more quickly. Unless otherwise stated, we used the same population genetic parameters as in Fig. 1.

**Figure S2.**
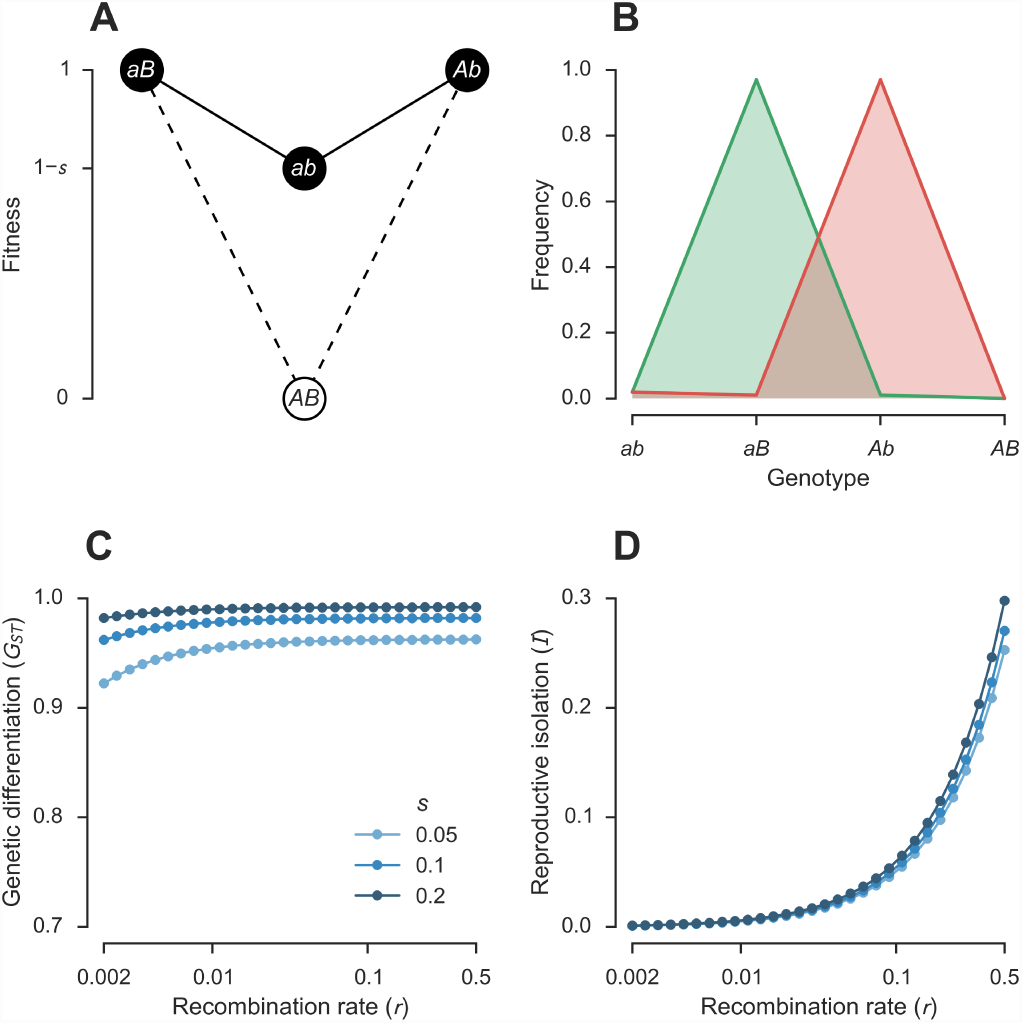
Selection for derived alleles in a single DMI can cause speciation. (A) Fitness landscape generated by a single DMI between *L* = 2 diallelic loci with selection for derived alleles. *s* measures the strength of selection against the *ab* genotype. This is an example of a “mutation-order” model [6] because it assumes that the different populations experience the same environment. Note that if *s* = 1, the *ab* genotype is lethal and the model reduces to a disconnected neutral network with two genotypes: *aB* and *Ab*. (B) Genotype frequencies at equilibrium for populations evolving under weak recombination and selection (*r* = 0.002, *s* = 0.05). There are two stable equilibria, one with 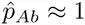 and the other with 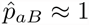 Both the genetic differentiation (C) and the reproductive isolation (D) among populations at the two stable equilibria increases with both *s* and *r*. The degree of reproductive isolation is well approximated by *I ≈ r*(1 + *s*)*/*2. Unless otherwise stated, we used the same population genetic parameters as in Fig. 1.

**Figure S3.**
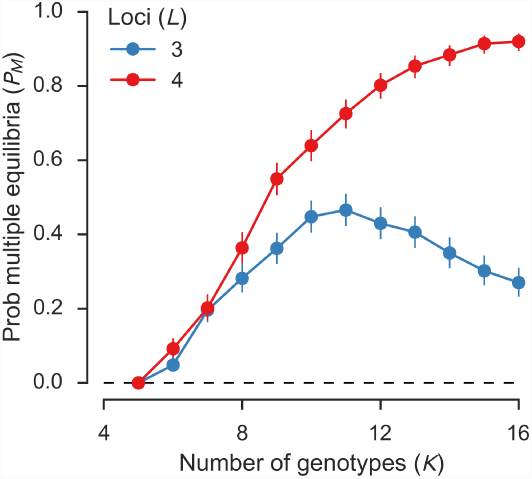
Speciation can emerge through the collective effects of multiple neutral DMIs that cannot, individually, cause speciation. Probabilities that neutral networks contain multiple stable equilibria (*P*_*M*_) based on ensembles of 500 random connected neutral networks for each combination of numbers of genotypes (*K*) and triallelic loci (*L*). See Fig. 2A for more details.

**Figure S4.**
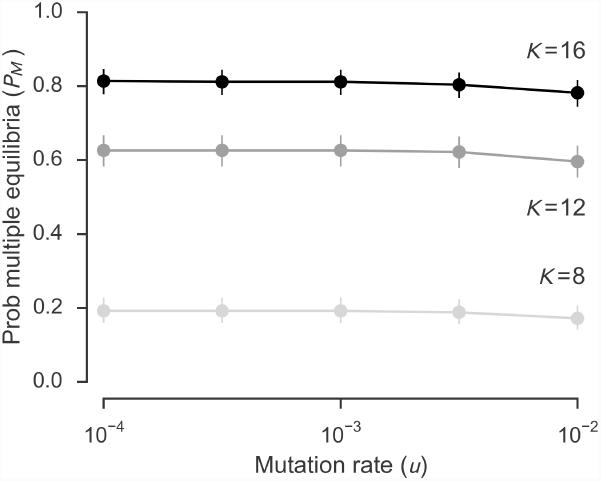
Changing both the mutation rate and the recombination rate while keeping the relative strength of recombination constant has only a small effect on emergent speciation. The three neutral network ensembles denoted by open circles in Fig. 2A were reanalysed for a range of mutation rates (*u*) and recombination rates (*r*) while keeping the relative strength of recombination constant (*r/u* = 4). See Fig. 2B for more details.

**Figure S5.**
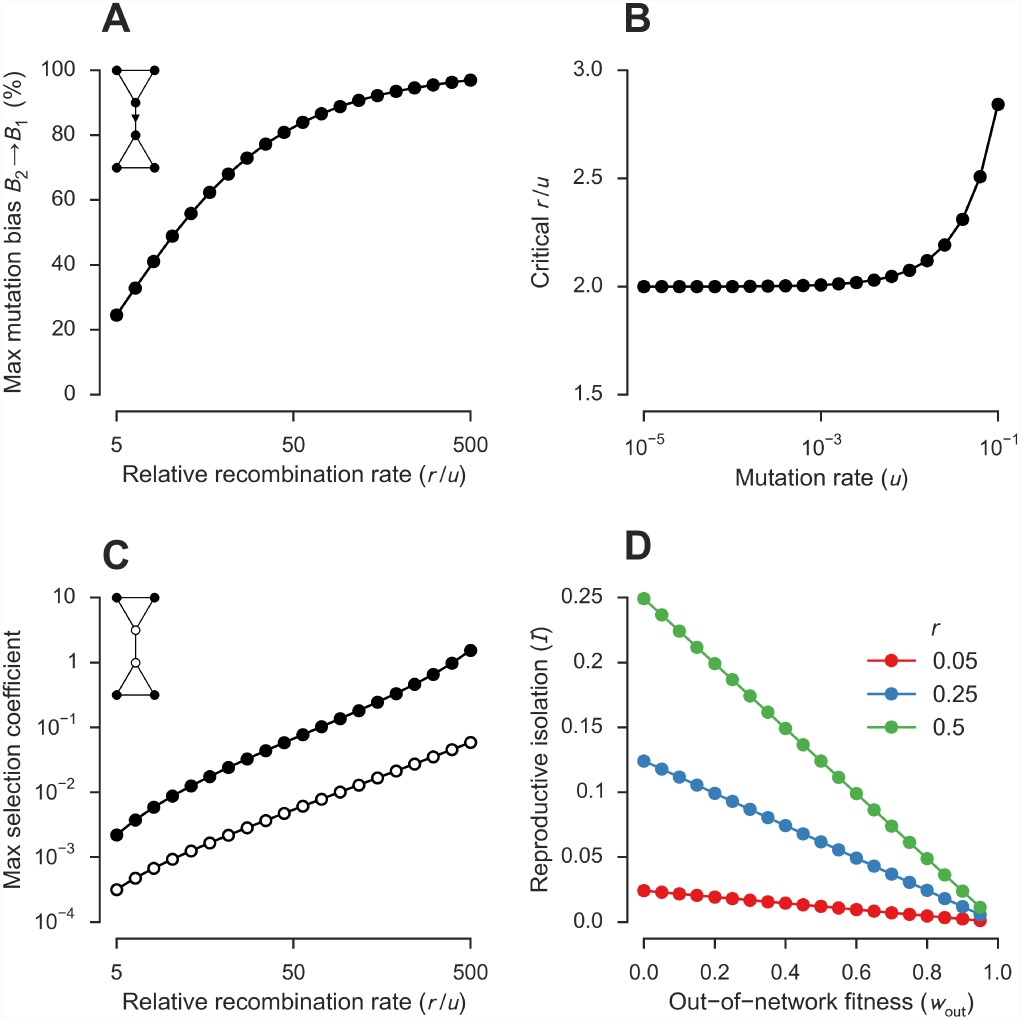
Emergent speciation is a robust mechanism. All analyses summarized here were carried out for the neutral network shown in Fig. 4A. Unless otherwise stated, we used the same population genetic parameters as in Fig. 2A. (A) Emergent speciation can occur in the presence of mutational biases. So far, all analyses have assumed that the mutation rates are equal for every locus in all directions. *B*_2_ → *B*_1_ mutations have the potential to perturb the *B*_2_ stable equilibrium. Values show the maximum increase in the mutation rate *B*_2_ → *B*_1_ (while keeping all other mutation rates constant) that allows the existence of two stable equilibria. For simplicity, we assumed that only one mutation was allowed per genotype per generation. The critical point at which the equilibria bifurcate (Fig. 4B) is approximately invariant with the relative strength of recombination (*r/u*). We calculated the critical recombination rate (*r*) for different values of the mutation rate *u* (spanning 4 orders of magnitude). We did this by finding the point where the symmetric equilibrium (in green in Fig. 4B and C) changes from stable to unstable. (C) Emergent speciation can occur in the presence of differences in the fitness of viable genotypes. The neutral network model assumes that all “in-network” genotypes have equal fitness, *w*. Values show the maximum selection coefficient, *s*_*i*_, by which the fitness of genotype *i* can be increased (*w*_*i*_ = 1 + *s*_*i*_), while allowing the existence of two stable equilibria. (D) Emergent speciation can occur in the presence of partial DMIs. The neutral network model assumes that “out-of-network” genotypes have a fitness of *w*_out_ = 0. Values show the effect of changing *w*_out_ on the degree of reproductive isolation among populations at the two stable equilibria: *I ≈ I*_0_(1− *w*_out_), where *I*_0_ *≈ r/*2 is the reproductive isolation when *w*_out_ = 0 (i.e., the default model). This is because higher values of *w*_out_ imply higher hybrid fitness.

**Figure S6.**
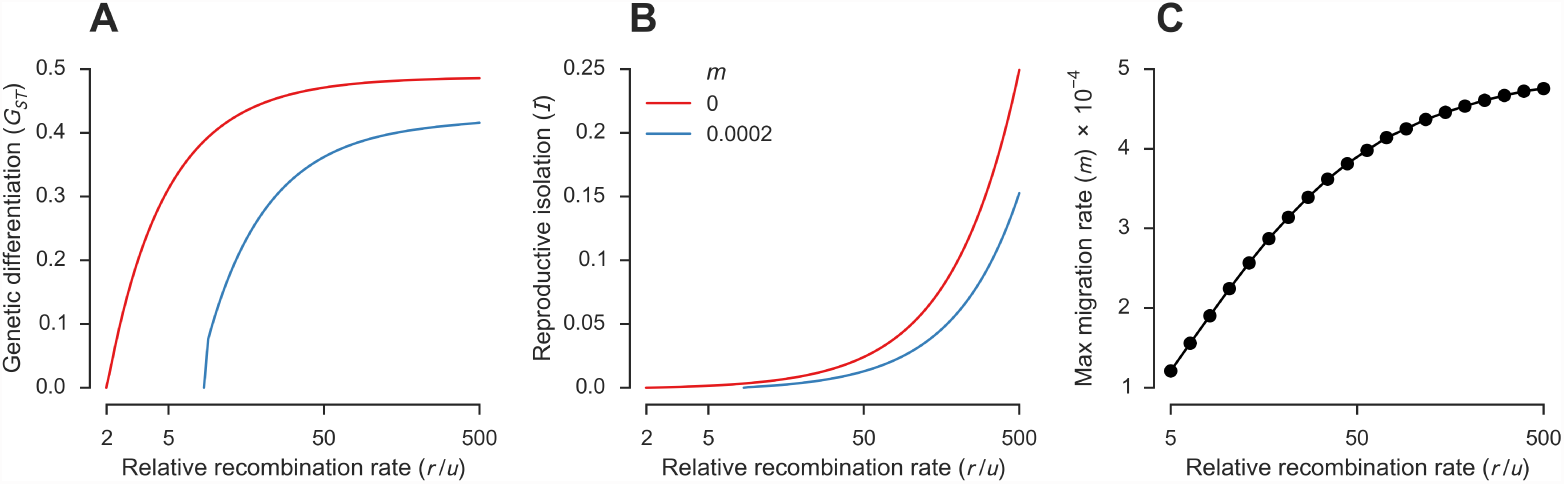
Emergent speciation can occur in the presence of gene flow between populations. Both the genetic differentiation (A) and the reproductive isolation (B) among populations at different stable equilibria for the network shown in Fig. 4A increases with the relative strength of recombination (red), and can persist in the presence of weak gene flow (blue). (C) Maximum migration rate, *m*, between two populations that allows the existence of two stable equilibria. The population genetic parameters were the same as those in Fig. 2A.

**Figure S7.**
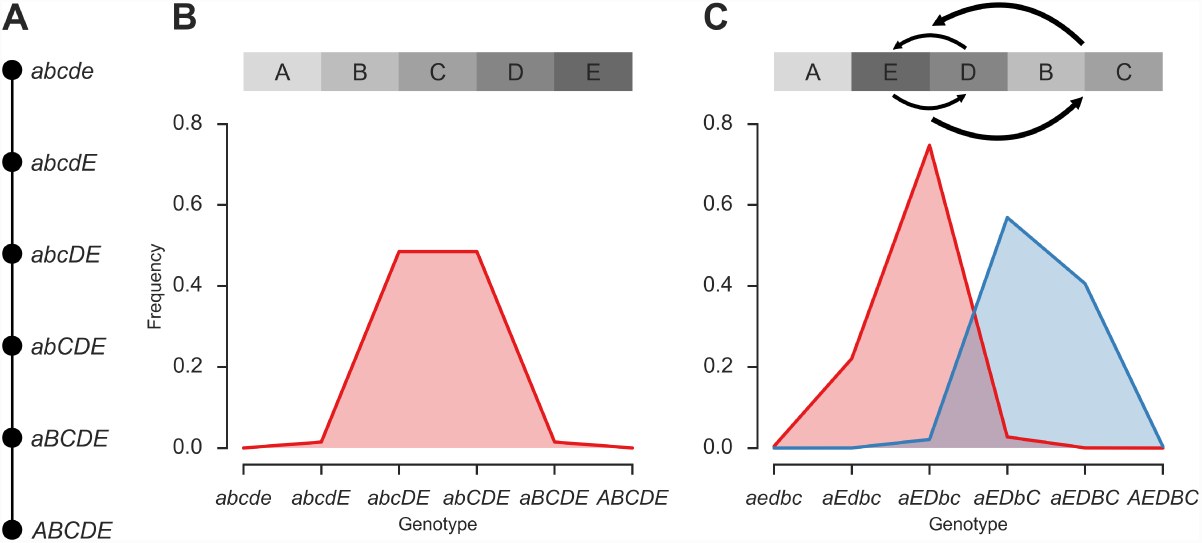
Emergent speciation depends on the precise pattern of recombination between sites and cannot be strictly predicted from the topology of a neutral network. (A) Neutral network of *K* = 6 genotypes generated by incompatibilities between *L* = 5 loci. (B) The neutral network shown in (A) shows a single stable equilibrium for any relative strength of recombination (0 *≤ r/u ≤* 500). The figure shows *r/u* = 64. (C) The genotype network derived from that shown in (A) by inverting the D and E loci and inserting them between the A and B loci has exactly the same topology as that in (A), but shows two stable equilibria when the relative strength of recombination is high (*r/u ≥* 3.65). The figure shows *r/u* = 64. The population genetic parameters were the same as those in Fig. 2A.

**Figure S8.**
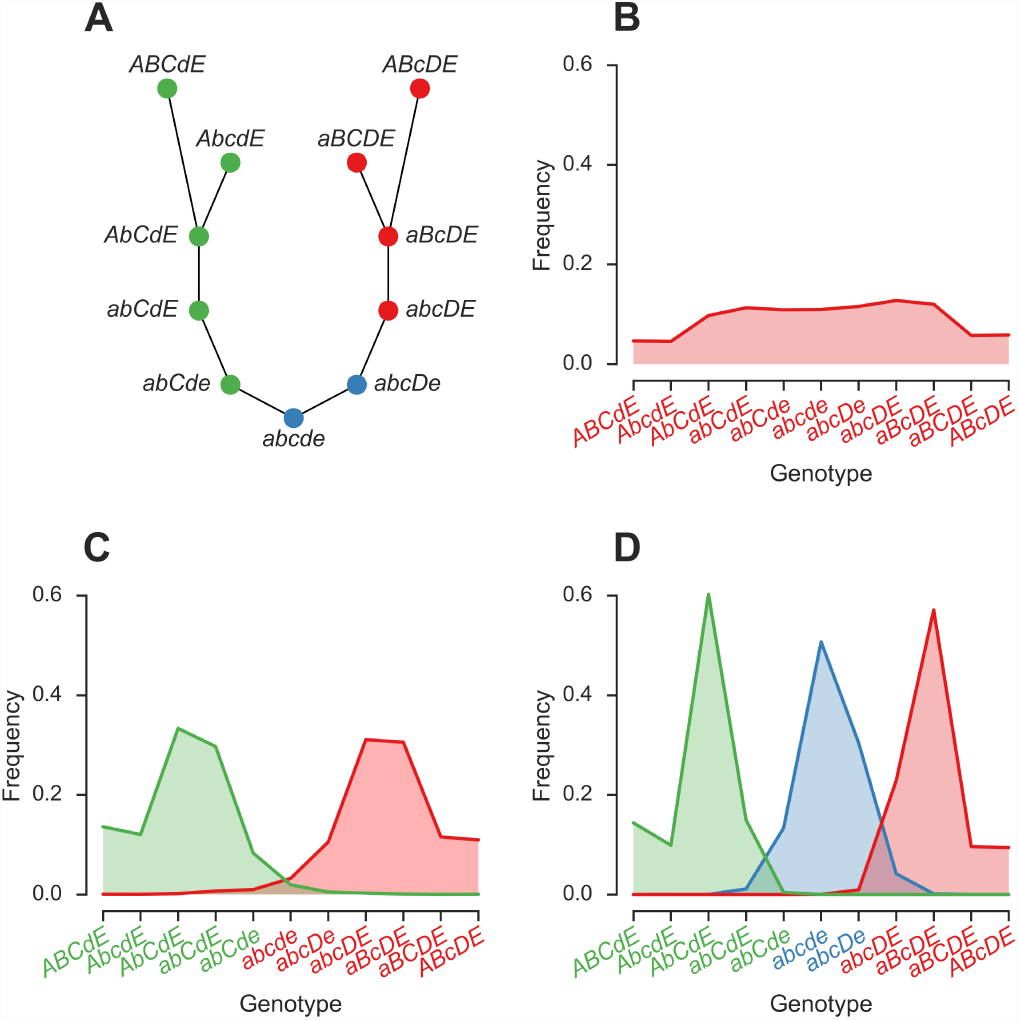
Example of a neutral network supporting emergent speciation. (A) Neutral network of *K* = 11 genotypes generated by multiple DMIs between *L* = 5 diallelic loci. Genotypes are represented by closed circles. Solid lines connect genotypes differing at a single locus. The colors relate to the equilibria as explained below. (B–D) The existence of multiple stable equilibria in the neutral network shown in (A) depends on the relative strength of recombination, *r/u*. Populations initially fixed for genotypes shown in red, green or blue, evolve to the equilibrium of the same color. (B) When the relative strength of recombination is low (*r/u <* 0.36), populations initially fixed for any of the 11 genotypes in the neutral network evolve to the same stable equilibrium. The figure shows the equilibrium when *r/u* = 0.1. (C) At higher relative strength of recombination (0.36 *< r/u <* 5.3), there are two stable equilibria. The figure shows the equilibria when *r/u* = 1. (D) At even higher relative strength of recombination (*r >* 5.3) there are three stable equilibria. The figure shows the equilibria when *r/u* = 10. The population genetic parameters were the same as those in Fig. 2A.

**Figure S9.**
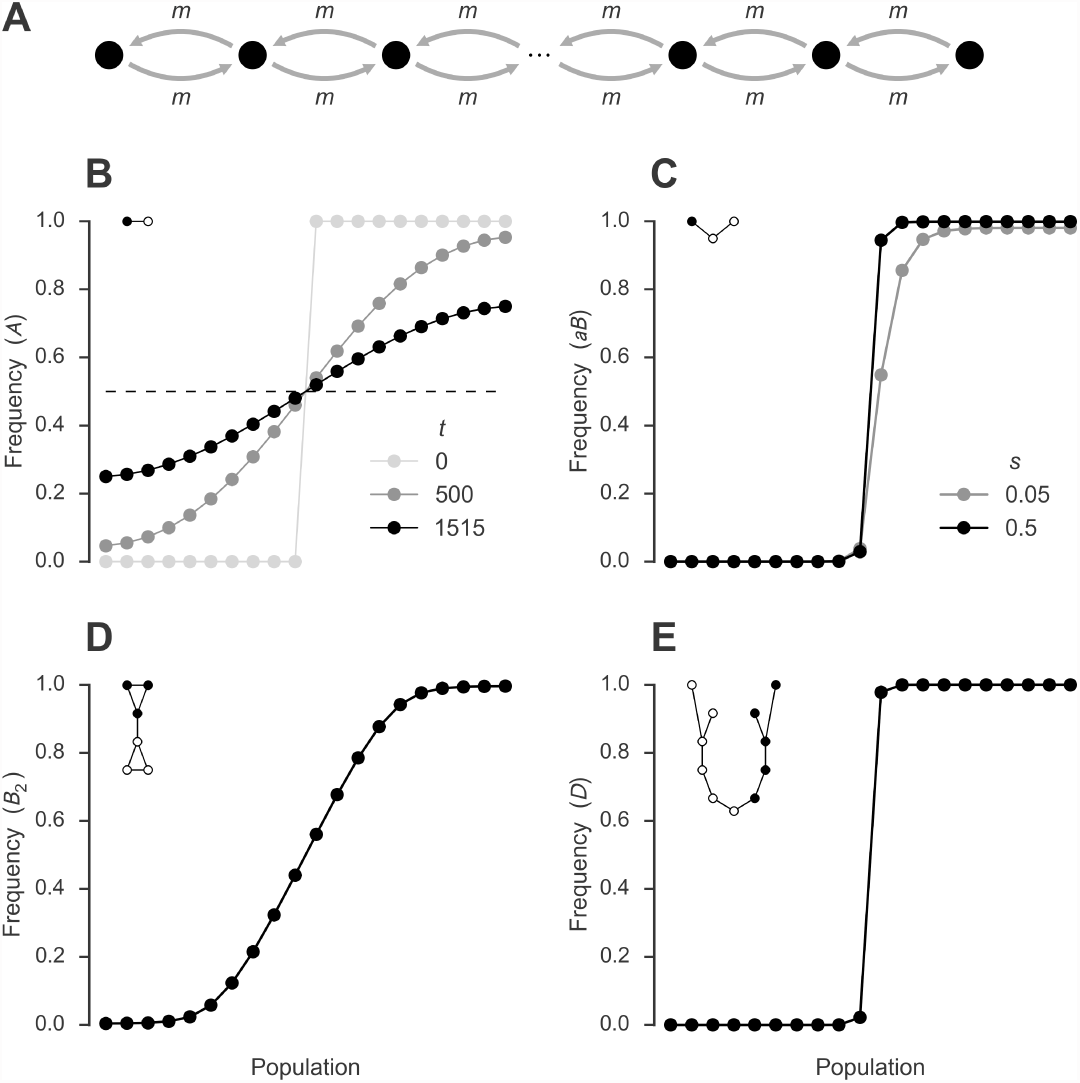
Neutral DMIs can generate strong reproductive barriers. (A) Stepping-stone model. A number *n* of populations are arranged in a line. Every generation a proportion 2*m* of a population emigrates to its two neighboring populations (except populations 1 and *n*, which have only one neighbor, so only *m* of each of them emigrate). (B) Spread of a neutral allele over *n* = 20 populations. Locus A has two neutral alleles, *A* and *a*. Initially, populations 1–10 are fixed for the *a* allele and populations 11–20 are fixed for the *A* allele (light gray). We allow the populations to evolve with *m* = 0.025, without mutation at the A locus. The dark gray points show the frequencies of the *A* allele after *t* = 500 generations. After *t* = 1515 generations, the frequency of *A* has increased from 0 to 25% in population 1 (black; *T*_*N*_ in Table 2). Eventually, the population will reach equilibrium with each neutral allele at a frequency of 50% in every population (dashed line). (C–E) Equilibrium allele or genotype frequencies in a hybrid zone formed after the contact of two populations initially at different stable equilibria on opposite ends of the line. See Table 2 for analysis of flow of unlinked neutral alleles through these hybrid zones. (C) DMI with selection (S2 Fig). Initially, populations 1–10 are fixed for the *Ab* genotype and populations 11–20 are fixed for the *aB* genotype. The points show the equilibrium frequencies of *aB* for *r* = 0.5 and *m* = 0.025 and different values of *s*. (D) Neutral network shown in Fig. 4A. Initially, populations 1–10 are fixed for the *B*_1_ allele and populations 11–20 are fixed for the *B*_2_ allele. The points show the equilibrium frequencies of the *B*_2_ allele for *r* = 0.5 and *m* = 0.025. (E) Neutral network shown in S8 Fig A. Initially, populations 1–10 are fixed for the *d* allele and populations 11–20 are fixed for the *D* allele (an indicator of the red equilibrium in S8 Fig). The points show the equilibrium frequencies of the *D* allele for *r* = 0.5 and *m* = 0.025. The population genetic parameters used in (C–E) were the same as those in Fig. 2A but with no mutation (*u* = 0) at the neutral marker locus and free recombination (*r* = 0.5) between all loci.

